# SAMPL-seq reveals micron-scale spatial hubs in the human gut microbiome

**DOI:** 10.1101/2024.10.08.617108

**Authors:** Miles Richardson, Shijie Zhao, Ravi U. Sheth, Liyuan Lin, Yiming Qu, Jeongchan Lee, Thomas Moody, Deirdre Ricaurte, Yiming Huang, Florencia Velez-Cortes, Guillaume Urtecho, Harris H. Wang

## Abstract

The local arrangement of microbes can profoundly impact community assembly, function, and stability. To date, little is known about the spatial organization of the human gut microbiome. Here, we describe a high-throughput and streamlined method, dubbed SAMPL-seq, that samples microbial composition of micron-scale sub-communities with split-and-pool barcoding to capture spatial colocalization in a complex consortium. SAMPL-seq analysis of the gut microbiome of healthy humans identified bacterial taxa pairs that consistently co-occurred both over time and across multiple individuals. These colocalized microbes organize into spatially distinct groups or “spatial hubs” dominated by *Bacteroideceae*, *Ruminococceae*, and *Lachnospiraceae* families. From a dietary perturbation using inulin, we observed reversible spatial rearrangement of the gut microbiome, where specific taxa form new local partnerships. Spatial metagenomics using SAMPL-seq can unlock new insights to improve the study of microbial communities.

**One Sentence Summary:** High throughput micron-scale subcommunity sampling and sequencing identifies distinct spatial associations of gut bacteria within and across individuals.

## INTRODUCTION

The human gut microbiome is stably colonized by hundreds to thousands of bacterial species^1^, which when perturbed has been associated with numerous diseases^2^. Beyond bulk compositional information, we know little about the micron-scale spatial assortment of microbes in the gut^3^. Microbes may spatially segregate due to metabolic and ecological interactions, ranging from cooperative sharing of niches to direct competition or antagonism^4^. As such, spatial organization can play a critical role in community makeup, function and stability^1,5^. In general, a spatially structured ecosystem better maintains species diversity than a homogenized microbiome^6^. Nutrients can further tune species interactions^7,8^. For example, dietary fibers are known to modulate short chain fatty acid (SCFA) production by bacterial consortia in the colon^9^. Mapping the local spatial arrangement of the human gut microbiome could reveal rules governing its organization, diversity, and resiliency in both healthy and diseased states.

Several high-resolution imaging-based approaches have been developed to map microbial spatial arrangements^10–15^. These methods, such as CLASI-FISH^10^, HIPR-FISH^12^, SHM-seq^15^, and SEER-FISH^14^, rely on highly-multiplexed barcoding and imaging setups to identify microbes in tissue sections. While these methods offer high spatial resolution and precise spatial coordinate information, they require prior metagenomic sequencing to obtain genomic information needed for probe design, need experimental validation of labeled bacterial taxa, and demand sophisticated imaging setups. Nevertheless, these approaches have been used to profile the human oral microbiome and the mouse gut microbiome with success^12,16,17^. However, the spatial organization of the human gut microbiome is more challenging to study due to its high taxonomic diversity and inter-personal heterogeneity and imaging-based strategies have not been applied to the human gut microbiome.

We previously described a spatial metagenomic sequencing approach (MaPS-seq^7^) based on analyzing “microbial plots”, which allows for the characterization of the bacteria present in hundreds of gut microbial sub-communities using metagenomic sequencing. However, the method required custom microfluidics, barcoded beads, and emulsion PCR steps that greatly limited throughput, scalability, and adoption. Recent single-cell sequencing advances in combinatorial split-and-pool barcoding (of beads for microfluidic encapsulation^18^ or single cells directly^19,20^) have streamlined the generation of large number of barcode combinations that significantly increased throughput and reduced cost/time. This combinatorial barcoding strategies could be adopted to label each “microbial plot” to achieve high-throughput sampling. However, these improvements have not been applied to complex microbial consortia, which would enable gut characterization at greatly increased scale to gain new insights previously not possible.

Here, we introduce **S**plit-**A**nd-pool **M**etagenomic **Pl**ot-sampling sequencing (SAMPL-seq), a streamlined spatial metagenomics method to analyze microbiome samples at micron-scale spatial resolution. By utilizing novel in-situ amplification steps to combine micron-scale particle-level spatial information with bacterial abundance, this is the first method to combine the community-sequencing approach of MaPS-seq with the high-throughput capacity of split pool barcoding. With these innovations, SAMPL-seq provides the necessary order of magnitude increase in scale and ease of use to enable in-depth spatial studies of the human gut microbiome. To demonstrate these new capabilities, we applied SAMPL-seq to human stool to reveal, for the first time, taxonomically distinct “spatial hubs” of the human gut microbiota that were stable over time and conserved between people. In response to dietary changes, these hubs reorganized into alternative spatial arrangements in a reversible manner, highlighting the flexible spatial assortment of the gut microbiome based on nutritional availability and environmental conditions.

## RESULTS

### Development of SAMPL-seq for microbial spatial metagenomics

SAMPL-seq utilizes the principle of microbial plot-sampling to identify bacteria that co-localize across tens of micrometer in natural sub-communities within a microbiome. An input microbiome sample (e.g., as little as ∼3 mm^3^) is first embedded and solidified in an acrylamide polymer matrix to preserve its original spatial organization (**Figure 1A**, **Methods**). This matrix contains acrydite linkers conjugated to a DNA adapter to facilitate downstream split-pool barcoding. The embedded sample is then cryo-fractured via bead-beating and the embedded bacteria are chemically lysed, while their DNA remains trapped in the gel. Next, the particles undergo three rounds of split-and-pool primer extension^18^ to create barcoded 16S rRNA primers that are unique to each particle and are filtered to a desired size (e.g., microbial plots of ∼40 μm in diameter, **Figure 1B**). An in-situ PCR reaction is performed to amplify the 16S rRNA V4 region across all particles using the now uniquely barcoded primers (**Supp. Figure 1A**). The PCR product is then UV-released from the particles, cleaned and concentrated (**Methods**). Sequencing and indexing adaptors are added by PCR and the library is sequenced on an Illumina platform. Reads thus contain both the 16S V4 sequence and a unique particle barcode (**Figure 1C,D**).

**Figure 1.**
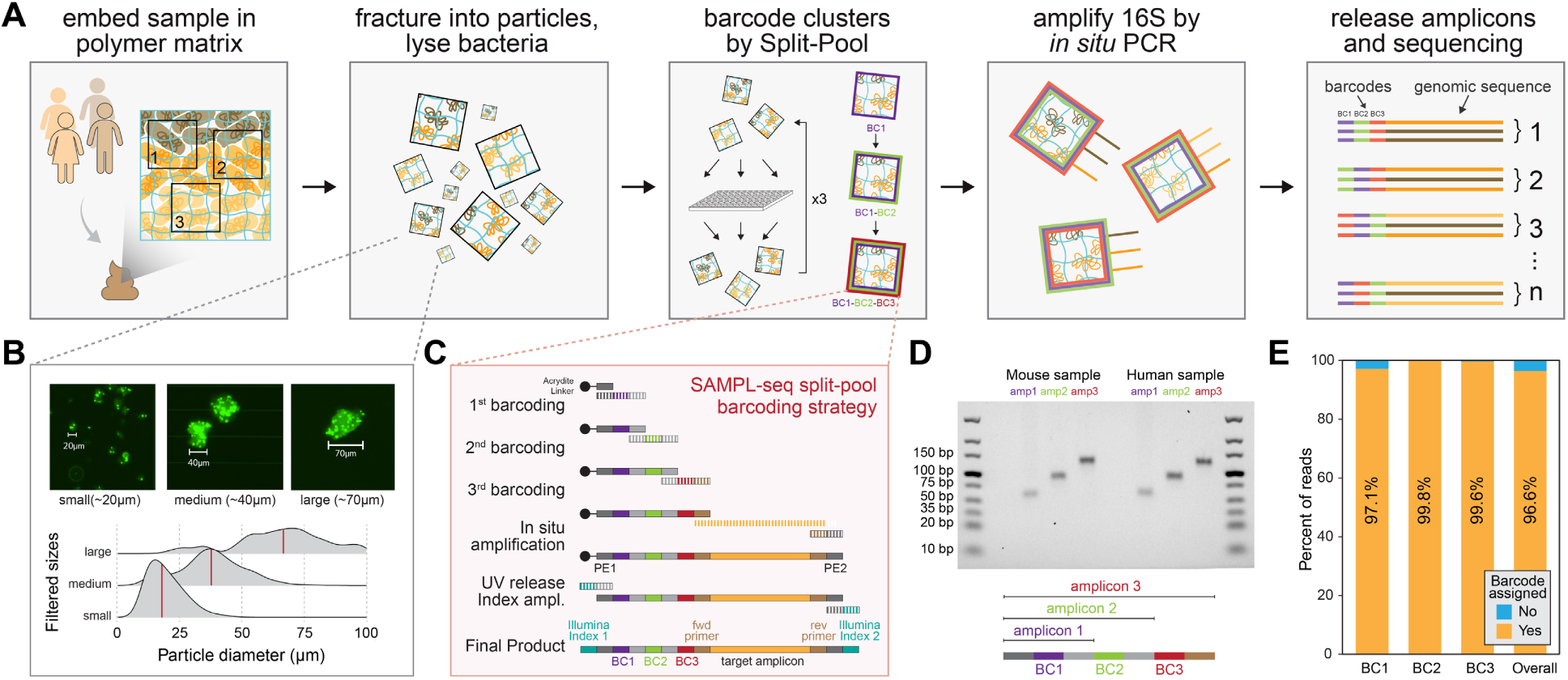
Spatial metagenomics of thousands of micron-sized communities using SAMPL-seq. **(A)** Step-by-step outline of the SAMPL-seq method**. (B)** Images of particles or “microbial plots” and their corresponding size distributions. **(C)** Schematic of split-pool barcoding steps that produce a barcoded primer, and downstream steps to generate final 16S amplicon library **(D)** Gel showing PCR products of fully extended barcodes from murine and human samples using primers that bind to different parts of the primer barcode sequence. **(E)** Barplot showing sequencing reads with successfully assigned barcodes across each barcoding step.

SAMPL-seq sequencing reads undergo barcode identification filtering with an overall success rate of ∼96.6% (**Methods, Figure 1E**). Reads are then grouped by particle according to their unique barcode combination, and amplicon sequence variants (ASVs, defined in this study as 100% sequence identical operational taxonomic units) are assigned using denoising. By replacing bead-based co-encapsulation described previously^7^ with in-situ split-pool barcoding and amplification, SAMPL-seq is substantially faster, scalable, and easier to implement to profile >10^4^ particles per sample without the need for microfluidics or other complex setups (**Suppl. Figure 1B, C**).

### Characterizing SAMPL-seq performance using mixed communities

We first characterized SAMPL-seq performance including replicability, overall bulk correlation, and throughput, along with spatial specificity, by determining background mixing rates of barcodes between particles. Two sets of mixing experiments, M1 and M2, were performed. In the M1 experiment, a homogenized microbiome sample (M1A) was mixed with a pure *S. pasteurii* culture (M1B) in two separate biological replicates (**Figure 2A, Suppl. Figure 2A, B**). Resulting SAMPL-seq data showed high experimental consistency between the M1A community in each replicate (*r* = 0.93, Pearson’s correlation) and high correlation with bulk 16S relative abundance (*r*=0.88, Pearson’s correlation) (**Figure 2B,C, Suppl. Figure 2C,D**). As expected, larger particle sizes tended to increase species diversity per particle (**Suppl. Figure 2E**). Further, the libraries had an overall multiplet rate^21^ of 4.7%, suggesting low mixing between communities (**Figure 2D,E**). Together, these results confirm that SAMPL-seq has high technical performance and reproducibility, good consistency with bulk sequencing results, and minimal methodological bias.

**Figure 2.**
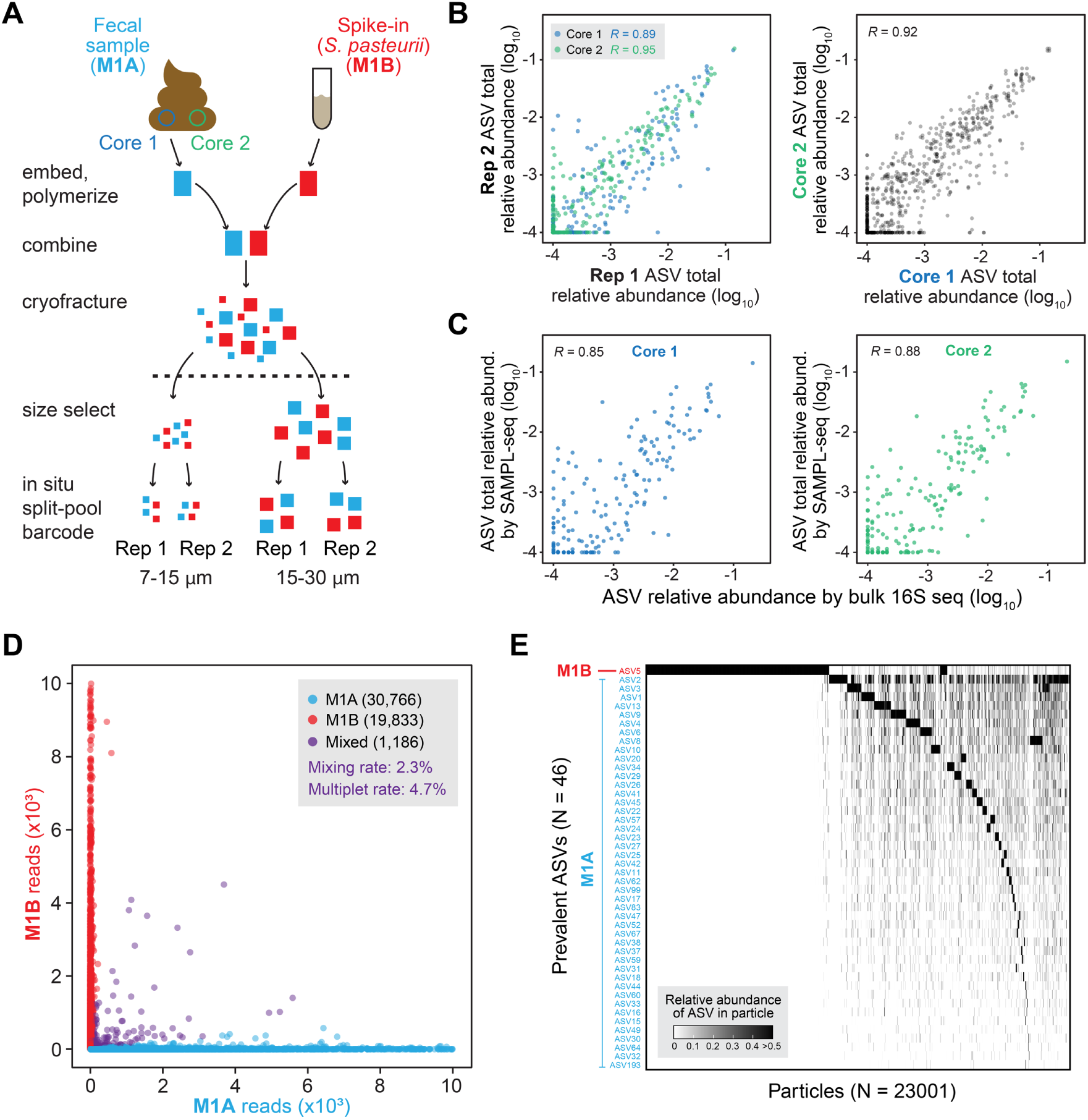
SAMPL-seq performance using mixing experiments. **(A)** An outline of mixing experiment M1. A human fecal sample and *S. pasteurii* are separately embedded and polymerized. They are then combined during the cryofracturing step, and are size sorted, and amplified and sequenced in aliquots of 10,000 particles. The homogenization procedure was repeated, for a total of two biological replicates. Two aliquots of 10,000 particles were sequenced as technical replicates. **(B)** Correlation of ASV relative abundance for technical replicates within each core and between cores. **(C)** Correlation of ASV relative abundance by SAMPL-seq versus bulk 16S sequencing for cores 1 and 2. **(D)** Scatterplot of particles showing the relationship between mixing and read count. **(E)** Heatmap of particles filtered to >50 reads per particle and prevalent (>1% RA) ASVs clustered by Bray-Curtis similarity and the Ward’s method.

In mixing experiment M2, two separate defined microbial sources of known composition were prepared at two cell densities of 2×10^8^ cells/μL (1x concentration) or 6×10^8^ cells/μL (3x concentration), separately embedded, and then mixed before cryo-fracturing (**Supp. Figure 3A**). The first source, M2A (Zymo D6331), consisted of common gut bacterial taxa at defined concentrations to allow for comparison to a known reference, while the second, M2B (Zymo D6320), consisted of a bacterial strain not present by design in M2A. After processing, each replicate yielded particles of mean size 50 μm in diameter (∼120-400 cells/particle) (**Supp. Figure 3B**). Reads from ∼16,000 particles across 5 replicates passed quality filtering (**Supp. Table 1**). Experimental replicates (1x versus 3x concentration) were highly correlated (*r*=0.84, Pearson’s correlation) (**Supp. Figure 3C**). The particle prevalence of each species, defined as percent of particles a species is found, also correlated well with its relative abundance as listed by the manufacturer (*r* = 0.80, Pearson’s correlation) (**Supp. Figure 3D**). Notably, at 3x bacterial input concentration, the species diversity per particle was higher (**Supp. Figure 3E**), which suggests SAMPL-seq’s sensitivity to different biomass levels. The average particle capture rate was 16.2% across replicates, which is on par with other single-cell methods^22^ (**Methods**). Importantly, only 1.4% of particles (177) contained mixed reads from both M2A and M2B sources. The overall multiplet rate, the mixing rate accounting for unobserved mixing, was 2.9% (**Supp. Figure 3F,G, Suppl. Table 2)**, which is also comparable to current split-pool methods^19,20,23^, with a low level of mixing between reads from different sources (**Suppl. Figure 3H**).

### Spatial metagenomics of the gut microbiome using stool material

Most microbiome studies rely on fecal matter as a reliable representation of the gut microbiome^24^. We sought to evaluate whether stool material can be used to assess the spatial architecture of the gut microbiome. SAMPL-seq was applied on three mouse gut compartments (small intestine, cecum, colon) along with the corresponding fecal pellets from the same mouse (**Suppl. Figure 4A**). ASV overall relative abundance and prevalence among particles were most similar between colon and stool than any other samples (r=0.55, p<2.2×10^-16^, r=0.71, p<2.2×10^-16^ respectively, Pearson’s correlation) (**Suppl. Figure 4B,C**). Consistent with our previous observations from the mouse gut microbiome^7^, the small intestine had a distinct set of spatially colocalized ASVs that persisted through the cecum and colon and remained colocalized in a subset of particles (**Suppl. Figure 4A**); this spatial signal could not be delineated from just bulk measurements. Principal coordinate analysis (PCoA) on SAMPL-seq particles from all compartments showed clustering between stool and colon, and clear separation from small intestine-derived samples (**Suppl. Figure 4D**). The cecum contained spatial signals from both small intestine and colonic communities. These results suggest possible spatial signals in stool samples that can be recovered with SAMPL-seq in a non-invasive manner to profile the *in vivo* colonic microbiome.

To explore the utility of SAMPL-seq for human gut microbiome studies, we applied the method to fresh stool from five healthy volunteers (H1, H10, H11, H18, H19), yielding data from >21,000 particles of ∼40 μm in diameter (**Suppl. Figure 5A-F**, **Suppl. Table 1, 3**). In one individual (H11), we performed additional longitudinal SAMPL-seq for five consecutive days (H11-D1 to D5) to explore temporal variation, yielding 18,000 particles. Unique ASV-particle barcode combinations saturated for detecting highly prevalent ASVs (>0.01%) and ASV-ASV co-occurrences, indicating sufficient sequencing coverage (**Suppl. Figure 5G-I**).Technical and biological SAMPL-seq replicates at Day 4 (H11-D4-R1 and R2) showed high correlation (*r*=0.92, p<2.2×10^-16^ and *r*=0.85, p<2.2×10^-16^ respectively, Pearson’s correlation), and longitudinal samples from H11 showed higher correlation than those from different donors (**Suppl. Figure 6A-E**). ASVs in the disrupted sample were consistently more prevalent across particles than in the original intact sample, showing that mechanical disruption eliminated the prior microbial spatial structure (**Suppl. Figure 6A**). The ASV abundance measured by SAMPL-seq and bulk 16S sequencing were highly correlated across all samples, indicating that taxonomic and compositional data was faithfully captured in these stool samples (**Suppl. Figure 6F,G**). While the microbiome composition was relatively consistent in H11 over 5 days (**Suppl. Figure 7A**), interpersonal samples exhibited greater compositional variation at the ASV level (**Suppl. Figure 7B**, *p* = 2.17 ×10^-5^ by Wilcox Rank-Sum test) than the family level (**Suppl. Figure 7C**). These results indicate that SAMPL-seq can be applied robustly to fecal samples despite natural variation in peoples’ microbiome, which allows further analysis of gut microbial spatial architecture.

### Identifying patterns of microbial spatial co-localization

To determine which ASV pairs are more or less likely to spatially localize in the human gut, we applied a null model based on a fixed-fixed permutation method, which is commonly used to find co-association patterns in ecological studies^25^ (**Methods, Figure 3A**). The model randomizes ASV presence across the dataset while preserving both the number of unique ASVs per particle and the prevalence of ASVs in the dataset. This model better accounts for the natural heterogeneity in particle-level ASV diversity compared to the Fisher’s exact test used previously^7^. With this null model, we could robustly detect the separation between M1A and M1B ASVs in the M1 mixing experiment, with minimal spurious associations (**Figure 3B**). Using this approach on temporal SAMPL-seq data (H11-D1 to H11-D5), we identified on average 86 statistically significant positive or negative co-associated ASV pairs in each day across a total of 73 ASVs (p<0.05, Benjamini-Hochberg (BH) false discovery rate (FDR)-corrected) (**Suppl. Table 4**). As a control, SAMPL-seq on a mechanically disrupted fecal aliquot of the H11-D4 sample showed substantially fewer co-associations (31 significant ASV pairs in disrupted versus 77 and 89 in intact Day 4 samples) and co-associations found in the disrupted sample had low correlation with the intact samples (**Suppl. Figure 8A,B**). Furthermore, we characterized the correlation in spatial associations of ASVs from three paired sets of fresh and frozen fecal samples and found high correlation between them (R = 0.88, 0.77, 0.80) (**Suppl. Figure 8C**). These results indicate that SAMPL-seq could be performed on frozen stool samples without the need for additional cryo-preservatives, which could allow retrospective analyses that leverage other existing stool biobanks^26^. Analysis across additional samples H1, H10, H18, H19 revealed striking patterns of pairwise ASV spatial co-associations. (**Figure 3C, Suppl. Figure 8D**)

**Figure 3.**
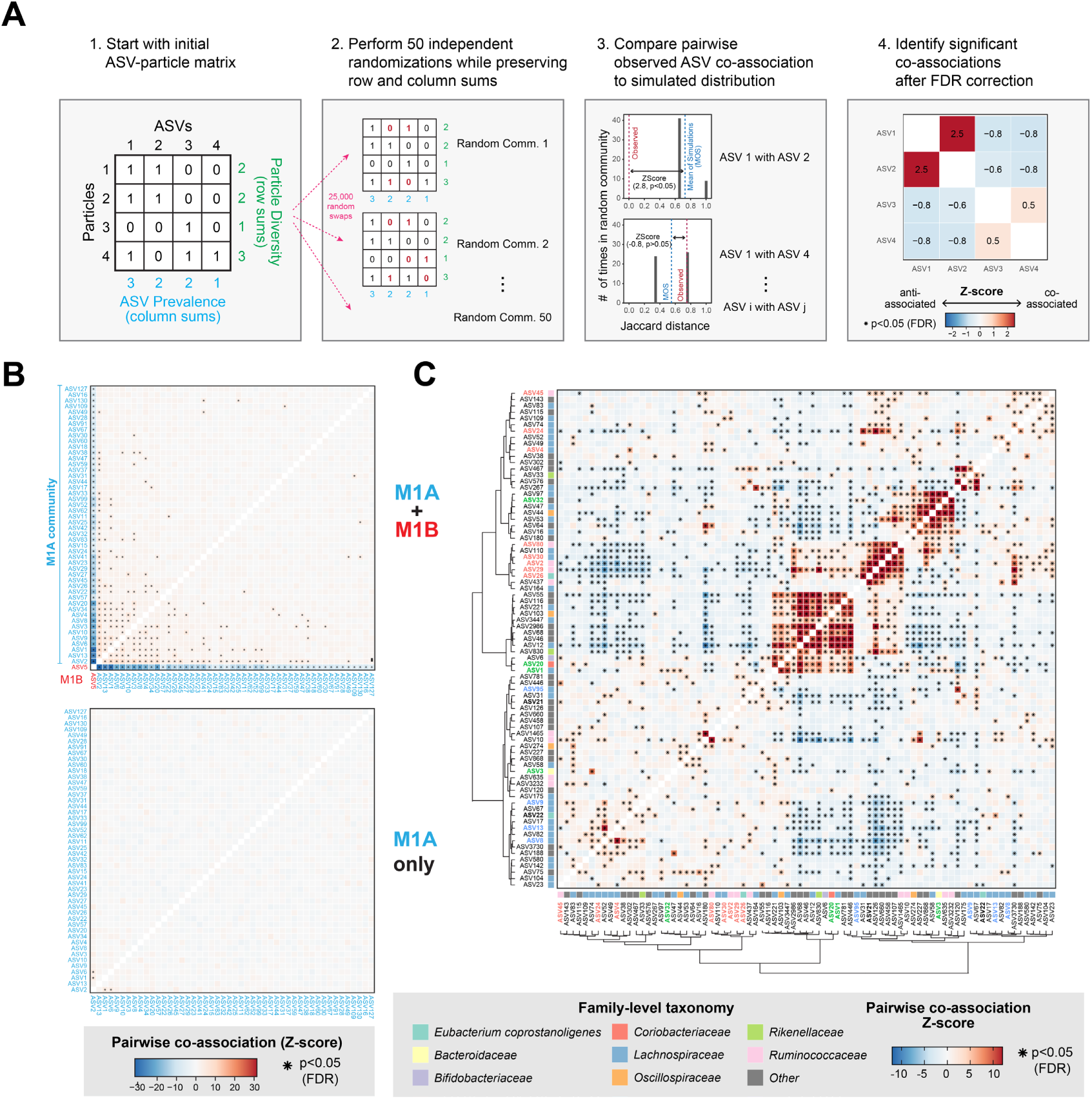
Colocalization analysis using SAMPL-seq data. **(A)** Diagram of the null model analysis. Particle count data is binarized and subjected to the *sim9* random swap algorithm. This is performed 50 times in parallel, and the resulting randomized data is used to generate a null distribution of co-associations between ASVs. Then, a Z-Score along with significance is calculated by comparing the observed co-association of an ASV pair to its null distribution. All pairwise associations are FDR corrected, and significantly co-associated pairs of ASVs are identified. **(B)** Pairwise co-association strength between ASVs in the Homogenized M1 mixing experiment. Stars correspond to statistical significance (p<0.05 FDR). Our method shows robust detection of the two separate communities M1A and M1B, along with minimal detection of significant associations in M1A alone. (**C**) Example of pairwise ASV co-association patterns in human sample H1 using the colocalization analysis.

Across the H11 longitudinal samples, we confirmed that the number of particles analyzed sufficiently captured the underlying spatial co-localization patterns. Our subsampling analysis shows that the number of subcommunities sequenced provide sufficient number to reach robust inference. Such inference requires at least thousands of particles^27^, which is only made possible with the throughput of our SAMPL-seq approach, which is superior to prior spatial metagenomic sampling methods (e.g., MaPS-seq) (**Figure 4A**). The spatial co-associations were consistent (i.e., 89.6% having same co- or anti-associations), indicating that a robust and stable spatial structure persisted over the 5-day sampling period (**Figure 4B, Suppl. Figure 8E-F**). To understand the overall spatial architecture in the longitudinal H11 dataset, we generated a co-association network using ASV pairs found across 2 or more days (**Figure 4C**, **Suppl. Table 5, Methods**). This spatial network of 33 ASVs could be grouped into four major clusters (L1-L4). Cluster L1 was composed of gram-positive *Ruminococcaceae* and *Lachnospiraceae*, with *Faecalibacterium prauznitzi* (ASV2) acting as a central hub that linked with all other ASVs in the cluster. In contrast, cluster L3 contained mostly *Lachnospiraceae* with a denser sub-network between *Fusicatenibacter saccharivorans* (ASV9), *Blautia massilliensis* (ASV13), *Blautia sp*. (ASV8), *Ruminococcus bromii* (ASV10)*, Dorea longicatena* (ASV16), and *Agathobacter rectalis* (ASV1). Another distinct cluster L2 contained mostly gram-negative *Bacteroidaceae* and *Parabacteroidaceae*, with *Bacteroides vulgatus/dorei* (ASV3) appearing as a central hub. *B. vulgatus* and *B. dore*i could not be uniquely resolved due to high 16S V4 similarity. Finally, cluster L4 contained *Eubacterium coprastanoligenes* (ASV22), *Alistipes marseille* (ASV27), and *Ruminococcus bicirculan*s*/champanellensis* (ASV4).

**Figure 4.**
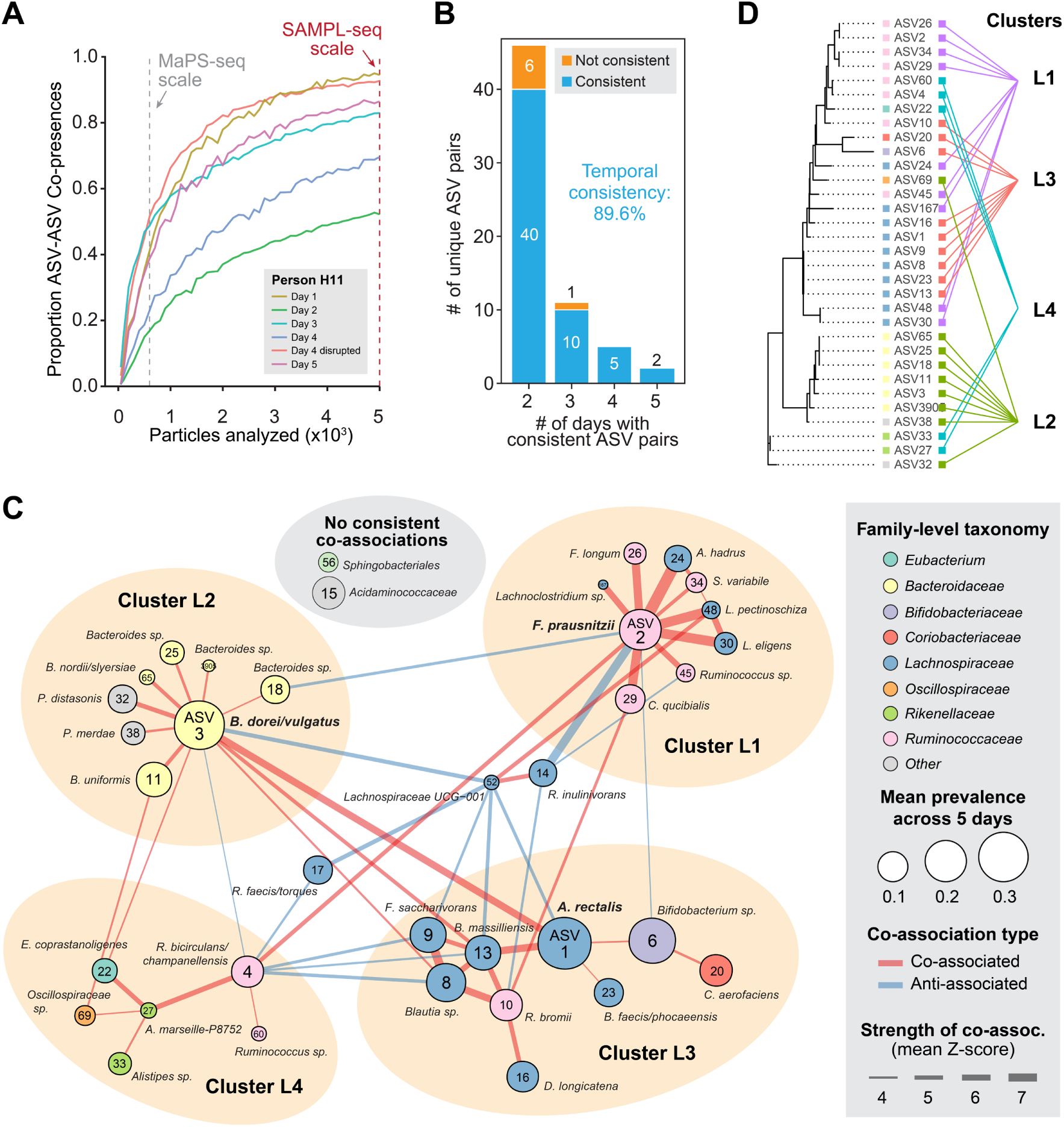
A longitudinal profile of gut microbiota spatial co-associations from one person. **(A)** Rarefaction plot of unique ASV-ASV co-occurrence in a particle (mean diameter ∼40 μm) (observed >3 times). SAMPL-seq scale shows the typical number of particles that could be sampled by SAMPL-seq, while MaPS-seq scale shows the typical number of particles that could be sampled by MaPS-seq. **(B)** Barplot of pairs of ASVs shared across days colored by the consistency of their association. (**C)** Network plot of ASV associations found on at least 2 days. Nodes are ASVs with size corresponding to mean prevalence across 5 days. Edges are associations strengths with color representing type. ASVs without edges did not have consistent associations across multiple days. (**D**) Phylogenetic tree of ASVs and their spatial hub cluster grouping.

Across clusters, a strong inter-phyla co-association was observed between *B. vulgatus/dore*i (ASV3) of L2 and *A. rectalis* (ASV1) of L3. Moreover, *R. bicirculan*s/*champanellensis* (ASV4) of L4 was co-associated with *F. prausnitzii* (ASV2) of L1 and anti-associated with several *Lachnospiraceae* from L3. To quantify the phylogenetic relatedness within spatial clusters, we calculated their respective net related indices (NRI), which showed clusters L2 and L3 individually having greater phylogenetic grouping than by chance (p=0.003 BH FDR-corrected, for both L2 and L3, **Suppl. Figure 9A**), and thus shared more similar phylogenetic assortment of ASVs (**Figure 4D**). Together, these results reveal a co-association network of temporally stable gut microbial assemblies that organizes into distinct “spatial hubs” with varying levels of phylogenetic relatedness.

### Conserved spatial hubs of gut microbiota across humans

We next sought to explore whether spatial co-association patterns were conserved across people. Even though many ASVs were unique to each person (i.e., only 12 prevalent ASVs were found in all 5 individuals), we identified a median of 261 significant co-associations across a median of 48 prevalent ASVs per individual. *F. prausnitzii* (ASV2) had the highest number of co-associations across the dataset (**Figure 5A**). Other *Ruminococcaceae*, including ASVs 29, 45, and 80, were also highly co-associated, while *Lachnospiraceae* ASVs 8, 9, and 13 were frequently anti-associated. ∼85% of ASV pairs had consistent co- or anti-associations in two or more people (**Figure 5B**, **Suppl. Fig 8E, Suppl. Table 6**). For ASV pairs found in three or more individuals, the spatial network showed three dominant hubs (P1-P3) (**Figure 5C**, **Suppl. Table 7**).

**Figure 5.**
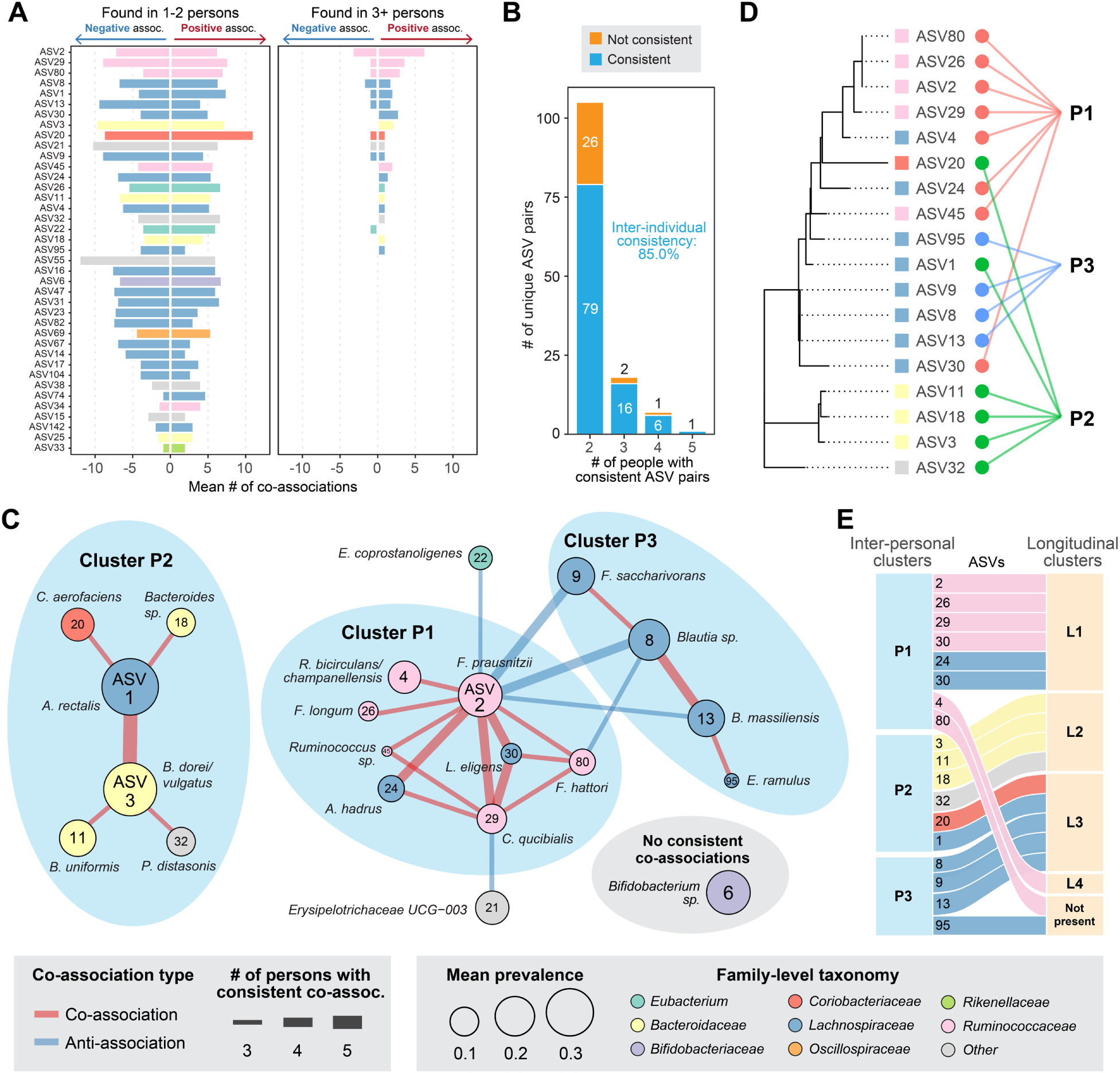
Consistent spatial hubs of gut microbiota found across humans. **(A)** Barplot of the mean number of co-associations for prevalent ASVs across all subjects (mean particle diameter ∼40 μm). **(B)** Barplot of pairs of ASVs shared across subjects colored by the consistency of their association. **(C)** Network plot of ASV associations found in least 3 subjects. ASVs without edges did not have consistent associations across multiple days. **(D)** Phylogenetic tree of interpersonal cluster members, along with taxonomy and cluster group. **(E)** Alluvial plot showing correspondence of ASVs from interpersonal spatial clusters (P1-P3) and longitudinal spatial clusters (L1-L4).

Hub P1 is highly connected, composed of *Ruminococceae* and *Lachnospiraceae*; *F. prausnitzii* was colocalized with all other cluster members (similarly to its hub architecture in L1), while *Cibiobacter qucibialis* (ASV29), *Lachnospira eligens* (ASV30), and *Faecalibacterium hattori* (ASV80) were also strongly co-associated Hub P2 contained *Bacteroides*, including *B. dorei/vulgatus* ASV3, along with *A. rectalis* (ASV1) and *Collinsella aerofaciens* (ASV20). The *B. dorei/vulgatus* and *A. rectalis* co-association was the strongest across both longitudinal and interpersonal datasets (L2 and P2 hubs). Finally, hub P3 is composed purely of *Lachnospiraceae*, including *F. sacchivorans* (ASV9), *Blautia massiliensis* (ASV13), and *Blautia sp* (ASV8). These P3 members were found to also co-associate in longitudinal cluster L3 in H11; they also showed strong anti-association with *F. prausnitzii* from hub P1, suggesting spatial segregation. Members of hubs P1 and P3 were significantly more related within each cluster than by chance (p=0.034, p=0.006 BH FDR-corrected) (**Figure 5D**, **Suppl. Figure 9B**).

Conserved longitudinal and interpersonal spatial patterns showed strong agreement, with ASV co-association pairs agreeing in their magnitude and sign (i.e., co- or anti-association) (Pearson’s *r*=0.7, **Suppl. Figure 9C, D**). The spatial grouping of ASVs in longitudinal (L1-L4) and interpersonal (P1-P3) hubs also showed significant overlap, as ASVs were more likely to be found in the same hubs than chance (Chi-Square Test, p = 0.001, **Suppl. Figure 9E**). The overlapping membership of longitudinal and interpersonal spatial hubs appears to be due to discrete sets of ASVs in both clusters; 6 ASVs present in P1 and L1, 4 ASVs present in P2 and L2, 3 ASVs from P3 and 2 ASVs from P2 forming L3 (**Figure 5E**). The taxonomic composition of our observed spatial groups is also noteworthy, with L2 and P2 dominated by *Bacteroides*, one of the core guilds in the microbiome^28^, while clusters L1, L3, P1, and P3 are dominated by *Firmicutes*, which belong to another main guild. Thus, spatial hubs present in both our interpersonal and longitudinal datasets indicate a consistent spatial pattern that is stable between people and over time and add to the evidence for conserved guilds in the human gut microbiome.

### Spatial changes of the human gut microbiome during a dietary perturbation

Diet can have profound impact on the gut microbiome both in terms of its composition and metabolism. However, we do not know how dietary changes alter the spatial arrangement of bacteria in the human gut. We therefore applied SAMPL-seq to uncover possible micron-scale changes in the spatial organization of the gut microbiome following a dietary intervention. We chose inulin as the perturbation since inulin is a common food component not metabolized by human enzymes, correlates with short-chain fatty acid fermentation, and can affect growth of beneficial commensal bacteria such as *Bifidobacterium*^29^. We gave individual H11 oral inulin supplementation (20 grams/day) in a 12-day study (Methods). Stool was obtained at baseline (4 days), during supplementation (4 days), and after discontinuation of supplementation (4 days). Both bulk 16S sequencing and SAMPL-seq were performed on these samples to assess compositional and spatial organizational changes.

Bulk 16S sequencing revealed no major alterations in the overall community structure (**Suppl. Figure 10**), consistent with previous observations^29^. With SAMPL-seq data, we first quantified the magnitude and total number of spatial interactions of an ASV by calculating its cumulative association Z-score (caZ-score) with all other ASVs, which showed large-scale spatial reorganization during inulin supplementation (**Figure 6A**, **Suppl. Figure 10**). While many ASVs had substantial caZ-score changes with inulin, including *F. prausnitzii* (ASV2) and *C. quicibialis* (ASV29), their abundance in the population did not change. This suggests that SAMPL-seq can identify alterations to the spatial organization of the microbiota that cannot be obtained via conventional bulk 16S analysis. For ASVs with the greatest overall changes in caZ-scores, we then assess their pairwise spatial co-associations (**Figure 6B**, **Suppl. Table 8**). With inulin, numerous ASVs had more spatial associations, suggesting the formation of new spatial pairings such as a notable triad of *L. pectinoschiza* (ASV48), *C. quicibialis* (ASV29), and *Lachnoclostridium sp*. (ASV167). When inulin is removed, these spatial structures also disappear, indicating an inulindependent change in the spatial organization.

**Figure 6.**
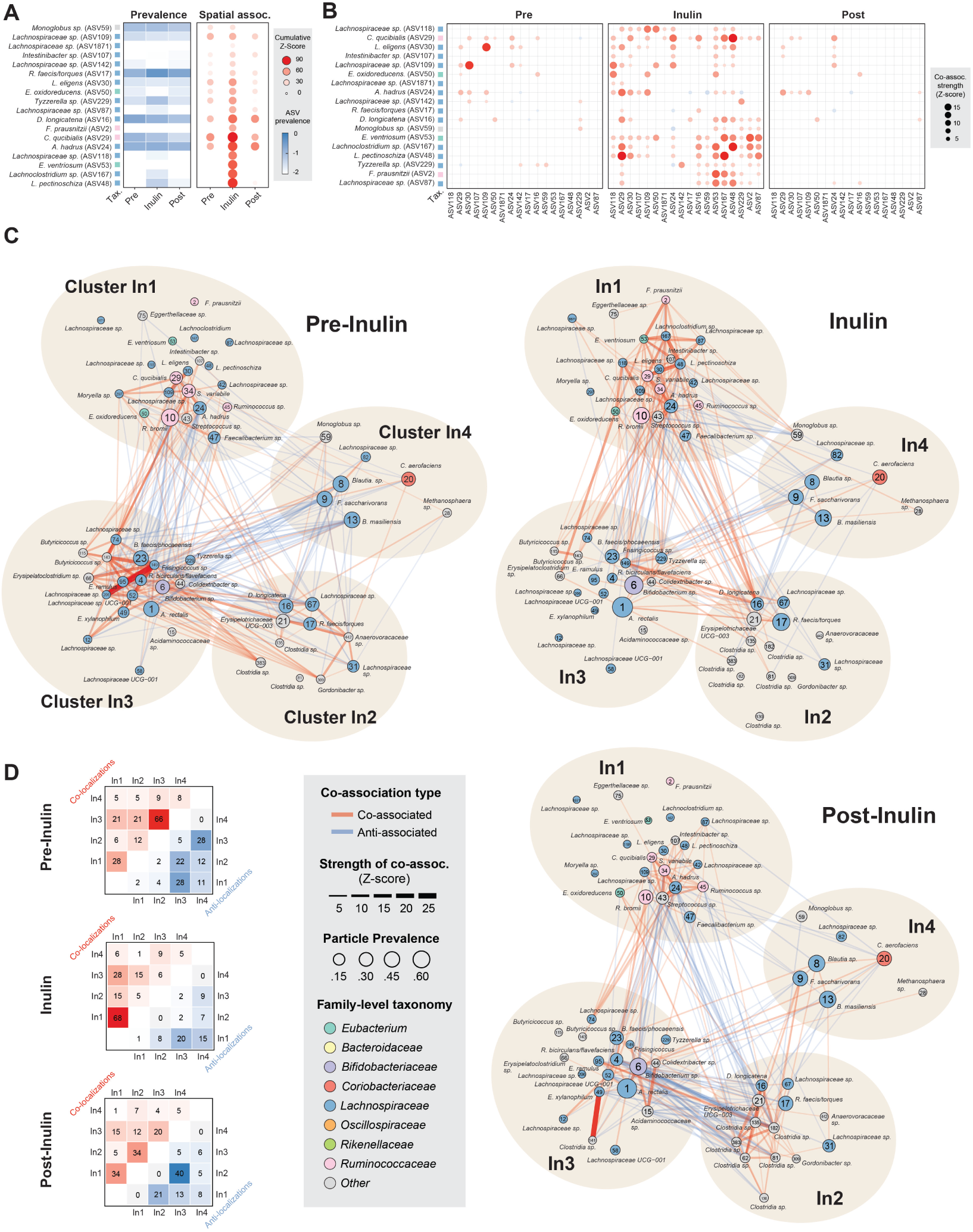
Spatial reorganization of the human gut microbiome in response to inulin supplementation. **(A)** Comparison of ASV prevalence versus cumulative spatial association for 16 ASVs that strongly respond to inulin supplementation (mean particle diameter ∼40 μm). **(B)** A dotplot of pairwise spatial associations among 16 inulin-responsive ASVs before, during, and post inulin supplementation. **(C)** Network plot of prevalent ASVs before, during, and post inulin supplementation showing 4 major spatial clusters (In1-In4). **(D)** Heatmaps summarize the number of positive (red) and negative (blue) spatial localizations found between ASVs within and between clusters.

We next visualized the entire co-association network to better understand the global spatial changes of all ASV pairs (**Figure 6C, Suppl. Table 9**). Four main inulin-mediated clusters emerged (In1, In2, In3, and In4), similar to the number of clusters previously in H11 (L1-4). Cluster In1, comprising of *Ruminococci* and *Lachnospiraceae*, shared substantial overlap with previously observed clusters L1 and P1. We then assessed the number of positive and negative co-associations within and across the four spatial hubs (**Figure 6D**). Prior to inulin exposure, Cluster In3 exhibited the largest number of within-cluster positive associations (66). With inulin, these In3 associations mostly disappeared (dropped to 6) while ln1 formed numerous new within-cluster spatial associations (totaling 68). When inulin was removed, within-In1 associations dropped back down to 34, but In3 associations did not fully recover to their pre-inulin levels (20 versus 66). Amongst these spatial changes, *L. pectinoschiza* (ASV48) was one of the major drivers of the observed spatial changes, with 20 new associations occurring only during inulin supplementation. *Lachnospira pectinoschiza* is an anaerobic gut bacteria known to utilize dietary fibers such as pectin^30^. Other key inulin-stimulated ASVs pairings in In1 involved *F. prausnitzii* (ASV2), *E. venturiosum* (ASV53), *C. quicibialis* (ASV29), and *Lachnoclostridium sp*. (ASV167) (**Figure 6C**), which aligns with previous documented evidence of inulin metabolism by members of the *Lachnospiraceae* and *Ruminococcaceae* families.^31^ Post-inulin, we found more spatial associations in In2 than before suggesting a spatial restructuring of the community. Nevertheless, the overall spatial patterns post-inulin was more similar to pre-inulin indicating reversible spatial restructuring by a dietary component. Collectively, these results highlight the coordinated spatial response of microbial hubs to the transient availability of a common dietary metabolite.

## DISCUSSION

Spatial metagenomics enabled by SAMPL-seq facilitates facile and high throughput delineation of microbial colocalization at the micron-scale. SAMPL-seq preserves spatial structure, as evidenced by low mixing rates, and allows the profiling of tens of thousands of “microbial plots” at a time, which is at least an order of magnitude improvement in scale over state-of-the art methods in plot sampling and key for accurate estimate of microbial co-localization.^27^ The ability of SAMPL-seq to provide high taxonomic resolution and local spatial information nicely complements imaging-based methods that can give global spatial positions of specific taxa^10–12,14,32^. Application of SAMPL-seq to stool samples yields microbiome co-association data that reflect the spatial organization found in the large intestine, thus allowing for non-invasive and longitudinal analysis of the colon at steady state and during dietary or other environmental perturbations.

Both the pairwise associations and spatial hubs found in the human gut may be hallmark features of a stable and healthy microbiota^33^, which when disturbed in disease states could lead to community-wide destabilization. Strains that grow together in spatial hubs may to be metabolically coupled or share a similar niche preference.^34^ Prior work suggests that *Bacteroides* form a dominant “guild” that are ecologically similar or metabolically complementary with one another in the Western adult gut,^28^ and our results showed that this group is also spatially organized as seen in clusters L2 and P2. *F. prausnitzii* is one of the most abundant butyrate-producing gut bacteria and its absence has been linked to disease-associated dysbiosis.^35^ The observed central role *F. prausnitzii* (ASV2) has in the P1 (and L1) spatial hub is particularly noteworthy, as it may indicate possible interspecies nutrient exchange. Indeed, past *in vitro* and *in vivo* experiments showed that *F. prausnitzii* grows better in the presence of other gut taxa.^36,37^ *Agathobacter rectalis* (ASV1) and *Bacteroides dorei/vulgatus* (ASV3) were observed as the most consistent and significant across all individuals. Both ASVs have been observed to localize in the mucus layer with *B. dorei* and *B. vulgatus* contributing to mucus degradation^38^ and *A. rectalis* showing preferential mucosal colonization despite an inability to utilize mucosal sugars^39,40^. Interestingly, *A. rectalis* can use sugars liberated by *Bacteroides* sp. to produce butyrate,^41^ which may explain their strong colocalization in the gut. Additional studies are needed to better elucidate the nature of these relationships *in vivo*.

During inulin supplementation, we observed notable changes in spatial associations including new spatial interactions between *L. pectinoschiza* (ASV48) and other *Ruminococci* including *F. prausnitzii* and *C. qucibialis*, which are involved in SCFA production and microbiome stabilization. Interestingly, *A. rectalis* (ASV1) and *Bifidobacteria sp.* (ASV 6), which are known to consume inulin^31^, did not form additional spatial co-associations during inulin supplementation, indicating a spatially-independent inulin metabolic process. Nevertheless inulin supplementation has been shown to enhance SCFA production^29^, which could be driven by the expanded co-associations within *Lachnospiraceae* and *Ruminococcaceae* families of cluster In1.

Further mechanistic experiments to probe the underpinnings that shape the observed microbial co-localizations could lead to better ways to modulate the gut microbiome and cultivate gut bacteria that have been recalcitrant to laboratory domestication.^37^ SAMPL-seq could be applied to other microbiomes such as those in soil or in foods to discover unseen spatially-mediated microbial interactions^42^ and build more accurate community-scale metabolic models^43^. With additional advancements, SAMPL-seq could evolve to encompass whole genome sequencing and incorporate genomic information from host cells, enabling us to associate spatial interactions with microbial genes, pathways, and microbiome-host spatial interactions.

## CONTRIBUTIONS

M.R., R.U.S. and H.H.W. conceived the project. R.U.S., M.R., and T.M. developed and validated the protocol. M.R., R.U.S., D.R., Y.H., L.L. J.L. and G.U. performed experiments. M.R., S.Z., Y.Q. and F.V.-C. analyzed the data. M.R. and H.H.W. generated and edited figures. H.H.W supervised the overall project. M.R., H.H.W, and S.Z. wrote the manuscript with input from co-authors. All authors reviewed and approved the manuscript.

## ACKNOWLEDGEMENTS

We thank Peter Sims, Dennis Vitkup, Lars Dietrich, Konstantine Tchourine for their immensely helpful insights and discussions, Georg Gerber and Travis Gibson for strengthening the computational rigor of the method, and members of the Wang Laboratory for providing a supportive environment. H.H.W. acknowledges funding support from the NSF (MCB-2025515), NIH (2R01AI132403, 1R01DK118044, 1R01EB031935, 1R21AI146817), ONR (N00014-18-1-2237), Burroughs Wellcome Fund (1016691), ARO (W911NF-22-2-0210), DARPA (HR0011-23-2-0001), Irma T. Hirschl Trust, and Schaefer Research Award. M.R., R.U.S. and F.V.C. were supported by the NSF Graduate Research Fellowship Program (DGE-1644869). R.U.S. was supported by the Fannie and John Hertz Foundation Fellowship.

## CODE AVAILABILITY

Scripts for read processing are implemented in BASH and R. They are available from https://github.com/wanglabcumc/SAMPL-seq

## DATA AVAILABILITY

Raw sequencing reads are available from PRJNA996899.

## METHODS

### Sample collection

Bulk human fecal samples were extracted from intact fecal sample using a sterile loop, placed in a cryovial, and stored at −80 C until use (IRB-AAAT4813). Samples used for strain isolation were extracted from an intact fecal sample using a sterile loop, and then added to sterile, pre-reduced PBS and processed in an anaerobic chamber. Sample were disrupted by vortexing, and then passed through a 40 μM filter. The resulting slurry was then diluted 1:1 with 50% glycerol in PBS, and stored at −80 C until use. SAMPL-seq human fecal cores were derived from intact fecal samples. Using the wide diameter end of P20 filter tip (Rainin), pieces of fecal sample were “cored”, and then immediately placed in tubes containing methacarn (60% methanol, 30% chloroform, 10% acetic acid). After 1 day of fixation, samples were removed from the P20 tip, and allowed to fix for an additional 12-24 hours. Then samples were transferred to 70% ethanol and stored at 4 C until use. Samples were used within one month. Mouse small intestine, cecum, large intestine and fecal samples were collected from a 12-week old Envigo Mouse (Protocol AABD4554). Samples were extracted and placed in methacarn for 24 hours. Once fixed, sections were cut to 3×3mm and used for downstream processing.

### Detailed SAMPL-seq protocol

#### Sample embedding

Fecal cores were cut to no larger than 3×3mm with a sterile razor to ensure full polymerization and placed in a sterile PCR tube. Disrupted fecal samples were generated by bead beating a 5mm diameter fecal pellet with 0.1mm glass beads for 1 minute at 4 C. Cores were then washed twice with 200 μL 1X PBS, then 200 μL permeabilization solution (1X PBS, 0.1% Triton-X 100 (vol/vol)) was added to the tubes, and samples were incubated for 5 min. Then, all excess solution liquid was removed from the tube, and samples were placed in a drying oven set to 90 C for 10 min. Once removed from the oven, samples were placed on ice to cool before embedding. The embedding solution contained 1x PBS, 10% (wt/wt) acrylamide, 0.25% (wt/wt) bisacrylamide, 5 μM primer (pe1) (**Suppl. Table 10**), 0.2% (wt/wt) 4-hydroxy-2,2,6,6-tetramethylpiperidin-1-oxy, 0.2%(wt/wt) tetramethylethylenediamine. The PE1 primer contains an acrydite group to enable adhesion to the gel, and a photocleavable spacer to allow for release using UV light. Samples were then covered with embedding solution to completely cover the sample (∼20-30 μL), and remained on ice for 5 min. The excess embedding solution was removed, and an additional fresh solution was added to cover the sample. Samples were then incubated on ice for 6-12 hours to ensure full perfusion. For final polymerization, excess embedding solution was removed, and samples were incubated on ice for 1 hour. After incubation, samples were placed in a 95C oven for up to 30min to ensure polymerization. Once embedded, samples were extracted from the PCR tube, excess polymer was trimmed using a sterile razor and washed with 1 mL PBS.

#### Particle fracturing

Samples were first placed in a stainless steel microvial (Biospec 2007). Next samples were frozen using liquid nitrogen for 2 minutes, without submerging the vial. Before proceeding, samples were shaken to ensure the sample could move freely. Next a single 6.35mm stainless steel bead (Biospec 11079635ss) was added to the vial, and the vial was plugged with a silicone rubber plug cap (Biospec 2008). Sample was then placed in liquid nitrogen for at least 2 minutes. Immediately the sample was transferred to a bead beater (Biospec 112011), and sample was beaten for 10 seconds at 3800rcf. Samples were then resuspended in 1 mL PBS. The suspended samples were then passed through a 100-micron cell strainer (Greiner Bio One 542100) into a new sterile tube. Particles were then washed twice more with 1X PBS. Washes were performed by spinning the sample down at 20,000 rcf, removing excess PBS without disturbing the particle pellet, and then adding 1 mL PBS.

#### Particle lysis

Particles were resuspended in 500 μL lysis buffer (Tris-HCl pH 8 10 mM, EDTA 1 mM, NaCl 100 mM), along with 375 U/μL lysozyme (Epicentre, R1810M), and incubated at 37C for 1 hour. Next samples resuspended in 500 μL digestion buffer (30 mM Tris HCl pH 8.0, EDTA 1 mM, 0.5% Triton X-100, 800 mM guanidine HCl) and 0.1ug/μL proteinase K (Epicentre MPRK092). Sample was then incubated at 65 C for 15 min, and then 95 C for 5 min to inactivate proteinase K. Particles were then washed three times with TET (10 mM Tris HCl pH 8.0, 1 mM EDTA, 0.1% Tween 20). If not proceeding to the next step, samples were brought to 15% glycerol, and frozen at −20C until further use.

#### Barcoding of particles via primer extension

This protocol uses a modified version of the procedures from Zilionis, et al^18^ to barcode primers present in the particles. The embedded primers in each particle are iteratively extended by primer extension over three rounds. All particle washes were done as follows: sample pellet was resuspended in 1 mL of washing solution, and then spun down at 20,000 rcf for one minute. The supernatant was removed. For each sample, a 96 well PCR plate was prepared with 1 μL of unique primer (**Suppl. Table 11)** distributed to each well. (pe1, pe2, pe3 primer sets). Samples were then washed 3 times with wash buffer (WB) (10 mM Tris HCl pH 8.0, 0.1 mM EDTA, 0.1% Tween 20), and adjusted to a volume of ∼833 μL. 110 μL 10X isothermal amplification buffer (NEB), 33 μL 10 mM dNTPs [0.3 mM final] (NEB), and 14 μL Bst2.0 8,000 U/mL [100 U/mL final] (NEB) was added to the sample. Then 9 μL particle/Bst2.0 mix was distributed to each well, either by pipet or using a Mantis liquid handler (Formulatrix). Plates were sealed and incubated at 60C for 30m. Then 20 μL of STOP25 (10 mM Tris HCl pH 8.0, 25 mM EDTA, 0.1% Tween 20, 100 mM KCl) was added to each well and plates were incubated at RT for 5 min. Then plates were pooled into a 5mL Eppendorf tube, and the total volume brought to 5mL with STOP25 to completely stop the reaction. The conical was then spun down at 20,000 rcf for 2 min, the supernatant was removed, and the pellet was transferred to a 1.5mL tube. The pellet was then washed 3 times with STOP10 (10 mM Tris HCl pH 8.0, 10 mM EDTA, 0.1% Tween 20, 100 mM KCl). To make ensure primers were single stranded for the next barcoding reaction, 1 mL freshly made DENATURE (0.5% Brij35, 150 mM NaOH) was used to resuspend the particles, and this was incubated at room temperature for 10 min. The particles were then washed three times with DENATURE, and washed once with NEUTRALIZE solution (100 mM Tris HCl pH 8.0, 10 mM EDTA, 0.1% Tween 20, 100 mM NaCl). This protocol was then repeated at the wash steps for each round of barcoding. If the protocol was stopped between barcoding rounds, the particles were washed three times with TET, brought to 10% glycerol, and frozen at −20C until continuing. Once barcoding was complete, incompletely extended primers needed to be removed. This is accomplished using hybridization to protect complete primers, and then Exo1 digest to remove the rest. Samples were washed 3 times with WB, and once with HYBRIDIZE (10 mM Tris HCl pH 8.0, 0.1 mM EDTA, 0.1% Tween-20, 330 mM KCl). Then the volume of the sample was adjusted to 300 μL with HYBRIDIZE, and 7.5μL of 1 mM 16S_515f_RC primer [∼20μM final] was added. This solution was incubated at 50C for 1hr to hybridize. Then, 50μL 10X ExoI buffer [1X final], 112.5 μL nuclease-free water, and 7.5 μL ExoI [0.3 U/μL final] were added, and incubated at 37 C for 1 hour. Then the tube was filled with STOP25, mixed, and incubated at RT for 5 min. This was then washed three times with STOP 10. Then it was incubated for 10 min at room temperature with DENATURE, and washed three times with DENATURE, once with NEUTRALIZE, and three times with TET, similar to the above barcoding. If stopped here, the solution was brought to 10% glycerol and stored at −20C.

#### Size filtering

To ensure consistent sizing, cell strainers were used. Samples were washed three times with PBS, and resuspended in 1 mL PBS. PBS was used as other buffers would impede flow through the filter. Samples were first passed through a 40 μM cell strainer (GBO 542140), and the strainer was washed with an additional 3mL PBS, and allowed to flow into the same tube. To recover particles larger than the filter, the strainer was inverted and placed onto a new tube. 1 mL of PBS was then passed through the strainer. This procedure was then repeated for using the smaller filtered fraction and a 20 μM cell strainer (GBO 542120). Once collected, all 5mL tubes were spun down at 20,000 rcf, the supernatant removed, and particles put into 1.5mL tubes. Then all samples were washed three times with TET and if proceeding, brought to 10% glycerol, and frozen at −20 C until continuing. Particles concentrations and sizes were determined by microscopy using a hemocytometer (Bulldog Bio DHC-N420). Particle were stained with SYBR green I (1x final) and imaged using a Nikon TI2 microscope. Particles were identified using the binary/define threshold function, and the equivalent diameter calculated using the NIS-Elements software.

#### In-situ PCR

Once quantified, particles were aliquoted for PCR reactions. Between 1000 and 10000 particles were amplified at a time. Particles were washed 3 times with TET, volume adjusted to 22.5 μL, and transferred to PCR tubes. PCR was then set up with the following reagents: 2.5 μL of 10μM pe2 816r REV primer [0.5μM final], 25 μL of KAPA Hifi 2X Readymix (Roche KK2601). It was then cycled with the following parameters: 98C 30s, 15 Cycles: 98C 10s, 55C 30s, 65 C 60s, extension 65 C 2min. Particles were then washed 3 times with TET. This was then repeated twice, for a total of 45 cycles.

#### UV release and magnetic bead cleanup

To release DNA from the particles, particle aliquots were washed 3 times with diffusion buffer (0.1% SDS, 1 mM EDTA, 500 mM ammonium acetate) and brought to a volume of 100 μL with diffusion buffer and transferred to PCR tubes. Particles were then placed on ice and treated with UV radiation for 15 min to break the photocleavable spacer. Aliquots were then incubated at 50C to allow for DNA diffusion into solution. Aliquots were then mixed at a 1:1 ratio with magnetic beads (Speedbeads Cytiva 65152105050250) and cleaned using a standard protocol. Cleaned DNA was eluted into 22 μL.

#### Indexing PCR

10 μL of purified PCR product was transferred to a new PCR tube, and the following reaction setup: 12.5 μL KAPA Hifi 2X Readymix (Roche KK2601), 2 mM SYTO9 (Thermo-Fisher, S34854), 1 μL forward index primer, 1 μL reverse index primer, 10.5 μL of in-situ PCR product. The samples were then run using a qPCR (BioRad CFX96) with the following program: 98C 45s, 30 Cycles 98C 10s, 68C 20s, 65 C 30s, Repeat, 65 C 120s, 10C Inf. Samples were removed during the extension phase if the appeared to leave the linear phase of PCR (usually between 14-20 cycles), and then replaced during the final extension. The resulting PCR product was assessed using a 2% acrylamide gel, the ∼490bp band extracted and purified (NEB Monarch, T1020L), and stored at −80 until use.

#### Sequencing and read processing

Samples were sequenced on the Nextseq 550 (Illumina) using the 150bp mid output or high output kit, depending on the number of samples, with a 30% phiX spike in. Over 10 million reads per particle library were targeted to ensure sufficient particle coverage after QC. The resulting sequencing reads needed additional processing to identify the particle barcode sequence. After demultiplexing, reads were demultiplexed using a custom BASH script (**Supplemental Materials**). Reads were first filtered using USEARCH 10 ^44^, with a cutoff of less than 1 expected error and minimum length of 150bp. Then, the particle barcode was identified and extracted from each read using ULTRAPLEX^45^ and a custom barcode mapping. 16S primers were then stripped, particle names were then modified using SeqKit^46^ to allow for recognition by USEARCH/VSEARCH ^47^, and reads of less than 69bp were removed and remaining reads were truncated to 69bp using seqkit. The result reads correspond to a 69bp 16S V4 region. Then, all samples except for the M2 mixing experiment were pooled together for denoising using UNOISE3^48^, and reads were mapped to ASVs using VSEARCH. For the M2 mixing experiment, reads were mapped directly to the reference 16S provided by the manufacturer. Since particle barcodes can contain errors, particle barcodes were extracted and subjected to error correction using the DNABarcodes^49^ package in R using an custom script (**Supplemental Materials**). Our barcode set allows for error correction of 1 base error, so barcodes with hamming distance larger than 1 were considered uncorrectable and removed. ∼96% of all barcode sequences were either correct or correctable. The resulting corrected ASV table was then used for subsequent analysis. For species level identification, 16S ASV sequences were matched to cultured strains from H1. If not present in the dataset, strains were matched to refseq 16S database at 100% identity. If no match was found, ASVs were labeled using the most specific taxonomic level available. The SINA Aligner ^50^ was used to create 16S rRNA alignments, which was then used to generate a phylogenetic tree with FastTree^51^. Taxonomy was also assigned with the SINA search and classify tool, and the SILVA^52^ taxonomy was used for downstream analysis.

### Detailed SAMPL-seq data analysis

#### Filtering

To remove potential read-through between particles, ASVs must be present at greater than 2% relative abundance within each particle to be considered “present”. Particles with less than 25 reads were removed from analysis. For visualization and co-association analysis, particles with 2 or fewer ASVs were removed, as it cannot be distinguished whether they represent a failed amplification or a monolithic community.

#### Rarefaction

Rarefaction was performed on individual amplification replicates, for the subset of ASVs > 1% particle prevalence across each amplification replicate. Unique ASV/particle pairs were used a the measurement as they represent “new” ASVs being found in new particles. Using reads from filtered particles, reads were sampled in 10 times at a given level, and the resulting number of unique read/particle pairs averaged at that point. This was repeated until the maximum number of reads was reached.

#### Co-localization

Co-association was quantified using a custom implementation of the SIM9 algorithm^25^, chosen for its low false positive rate, as implemented in the “sim9_single” function in EcoSimR package. The script used along with an example are included (**Supplementary Materials**). In brief, on each set of particles from one individual, a binarized (presence-absence) ASV table is subjected to a random swap, which preserves the ASV prevalence and particle diversities. Since this step only swaps a subset of values, it is performed 25,000 times to generate a “randomized” community based on the original diversity of the dataset. 50 of these randomized communities are generated to generate a null distribution of ASV co-localization. Then, the observed co-localization is compared to the distribution using a Z-test, and the resulting significance is subjected to FDR correction using the Benjamini-Hochberg procedure, with significance being an FDR-corrected p-value <0.05.

#### ASV Association Networks

The longitudinal association graph was generated by subsetting to ASVs pairs found to be significantly associating on 2+ days, and averaging the Z-score over that time. The interpersonal association graph was generated by subsetting to ASVs pairs found to be significantly associating in 3+ donors, and averaging the Z-score over that time. ASVs in each graph was then clustered using the spinglass clustering method as implemented in igraph^53^.

#### Net Relatedness

Net relatedness was calculated using the function “ses.mpd” from the R package *Picante*^54^. The taxa labels of each cluster were randomized 10,000 times, and the random MPD distribution was used to calculate the p-value. The p-values were then corrected using the Benjamini-Hochberg FDR correction.

#### Interpersonal Distance at the ASV or Family Level

Bray-Curtis distances were calculated between individual donors, using either ASV relative abundances or aggregated family-level relative abundances, and compared using a Wilcox Rank-Sum test to determine if distances were significantly higher when looking at the family level.

#### Plotting

Plotting for most graphs was performed using ggplot2^55^. Correlation was added to plots using ggpubR^56^. Statistical tests were performed using R 4.0. ASV association graphs were generated using ggraph^57^. Particle level heatmaps were generated using the geom_tile() function in ggplot. Particles were clustered using the Simpson overlap at the sample level, and ASVs were clustered by their Jaccard overlap across all particles in the heatmap.

#### Barcoding validation experiment

To validate the presence of barcodes after barcoding, aliquots of 10,000 barcoded but unamplified particles were aliquoted and subject to UV release, as described above. Then, the purified DNA was subjected to PCR using primers targeting anchor regions of the primers, with primer PE1 serving as the forward primer. For the reverse primers, Anchor 1-RC targeted the first extension, Anchor 2-RC the second, and 515RC the full length of the primer (**Suppl. Table 8**). Reactions were setup with 5 μL KAPA Hifi 2X Readymix (Roche KK2601), 1 μL forward primer (.3μM), 1 μL reverse primer (.3μM), 2 μL of cleaned primer DNA, and 1 μL of nuclease free water. Cycling was performed at 98C 3min, 30 Cycles 98C 20s, 60C 20s, 65 C 20s, Repeat, 65 C 120s.

#### Mixing experiments

Mixing rate calculations were performed two ways. In the first case, two bacterial communities were assembled: a homogenized fecal sample, and a pure culture of *Sporosarcina pasteurii*, and environmental bacteria not found in the gut. The homogenized fecal community M1A was generated by bead beating a 5mm diameter fecal pellet with 0.1mm glass beads for 1 minute at 4 C. The resulting solution was passed through a 40 μm cell strainer. Each was fixed in methacarn, washed with PBS, and subjected to the same embedding and polymerization protocol described above. Once polymerized and washed, samples were then subjected to cryo fracturing together in replicate with equal volumes of each polymerized community. Once co-fractured, the mixed community was treated as a single community and subjected to the rest of the protocol described above. In the second case, a mixed community was generated using two defined communities, the ZymoBIOMICS Gut Microbiome Standard (Zymo D6331) and ZymoBIOMICS Spike-in Control I (D6320). Cell concentrations were matched between them (2 x 10^6^ and 6 x 10^6^ cells per μL). Each was embedded separately in equal volumes of embedding solution. As above, equal volumes were mixed during cryo-fracturing, and were processed according to the protocol as described above. The mixing rate was calculated using the percentage of the particle assigned to the spike in community, either the *S. pasteurii* or ZymoBIOMICS Spike-in Control. Particles were considered mixed if they contained between 10-90% of the spike in. The multiplet rate was calculated as implemented as previously described^21^. The particle capture rate was calculated by dividing the number of particles after QC per library by either 10,000 or the number of particles identified before QC, whichever was greater. 10,000 was the estimated number of particles added to the sample for sequencing quantified by hemocytometer.

#### Bulk 16S sequencing

Bulk 16S samples were acquired by chemical or physical lysis. For chemical lysis, 3×3mm fecal samples were washed twice with PBS. Then sample was homogenized by vortexing in 500μL lysis buffer (10 mM Tris-HCl pH 8, 1 mM EDTA, 100 mM NaCl). Then lysozyme was added to the sample (final concentration ∼375 U/μL). The sample was vortexed and then incubated at 37 C for 1hr. Then 500 μL digestion buffer (50 mM Tris HCl pH 8. 0, 1 mM EDTA, 1% Triton X-100, 1600 mM guanidine HCl) was added along with proteinase K to 0.1ug/μL. Sample was vortexed again, and placed at 65 C for 15 min. Then, 100 μL of lysate was removed and subjected to a 1X bead cleanup, and resuspended in 22 μL of nuclease free H2O. Physical lysis was performed using our established sequencing pipeline, without spike-in^58^. Dual indexing amplification was performed using a modified protocol^59^. TruSeq 16S versions of the Earth Microbiome 515F and 806R^60^ matching those used the SAMPL-seq protocol were used for the first round of amplification, and standard TruSeq indices were used for the second round of amplification. Both rounds were performed using a qPCR, with samples removed before the end of linear amplification, usually between 8-12 cycles. Bulk samples were then pooled with SAMPL-seq libraries for sequencing.

#### Frozen Sample Processing

Fecal sample cores were taken from intact fecal samples, as described earlier. These cores were divided in half, and one half fixed immediately in methacarn at RT and processed as described earlier. The other half was immediately placed in a −80C freezer and kept frozen for up to 1 week. When ready for SAMPL-seq processing, the frozen sample was placed into pre-chilled (−20C) methacarn, and fixation proceeded at 4C for 24 hrs. Once fixed, the sample was processed as described earlier.

---

**Supplementary Figure 1.**
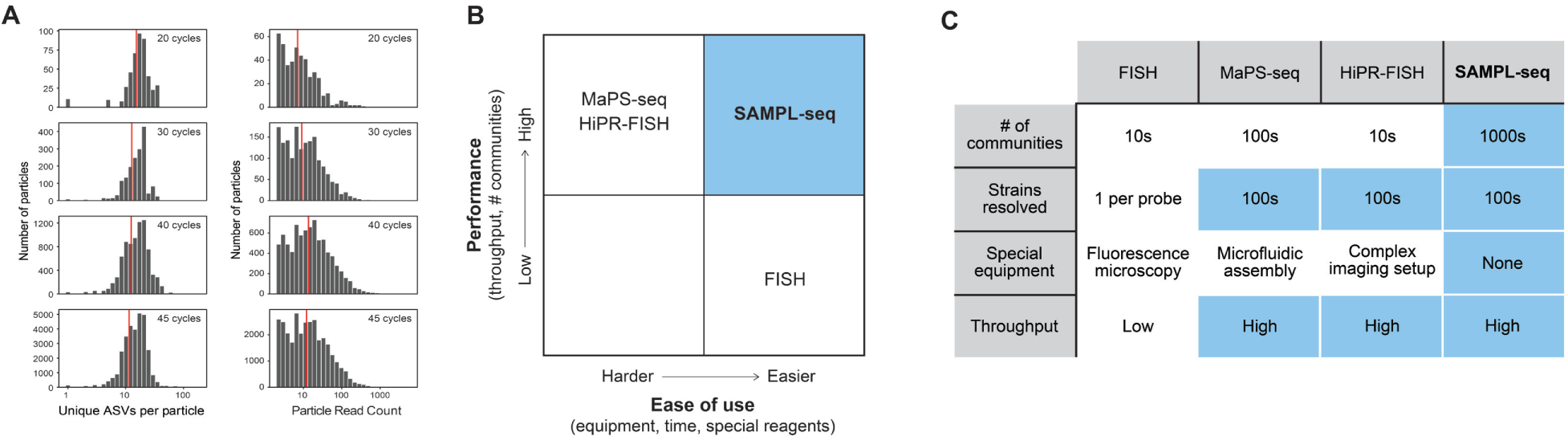
Characterization of SAMPL-seq steps and comparison with other methods. **(A)** Histograms showing the effect of the number of in situ PCR cycles on both the ASVs per particle and reads per particle based on different number of PCR cycles. **(B)** Plot summarizing the overall ease of use and performance of various microbial spatial analysis methods. **(C)** Table comparing the performance of different spatial analysis methods.

**Supplementary Figure 2.**
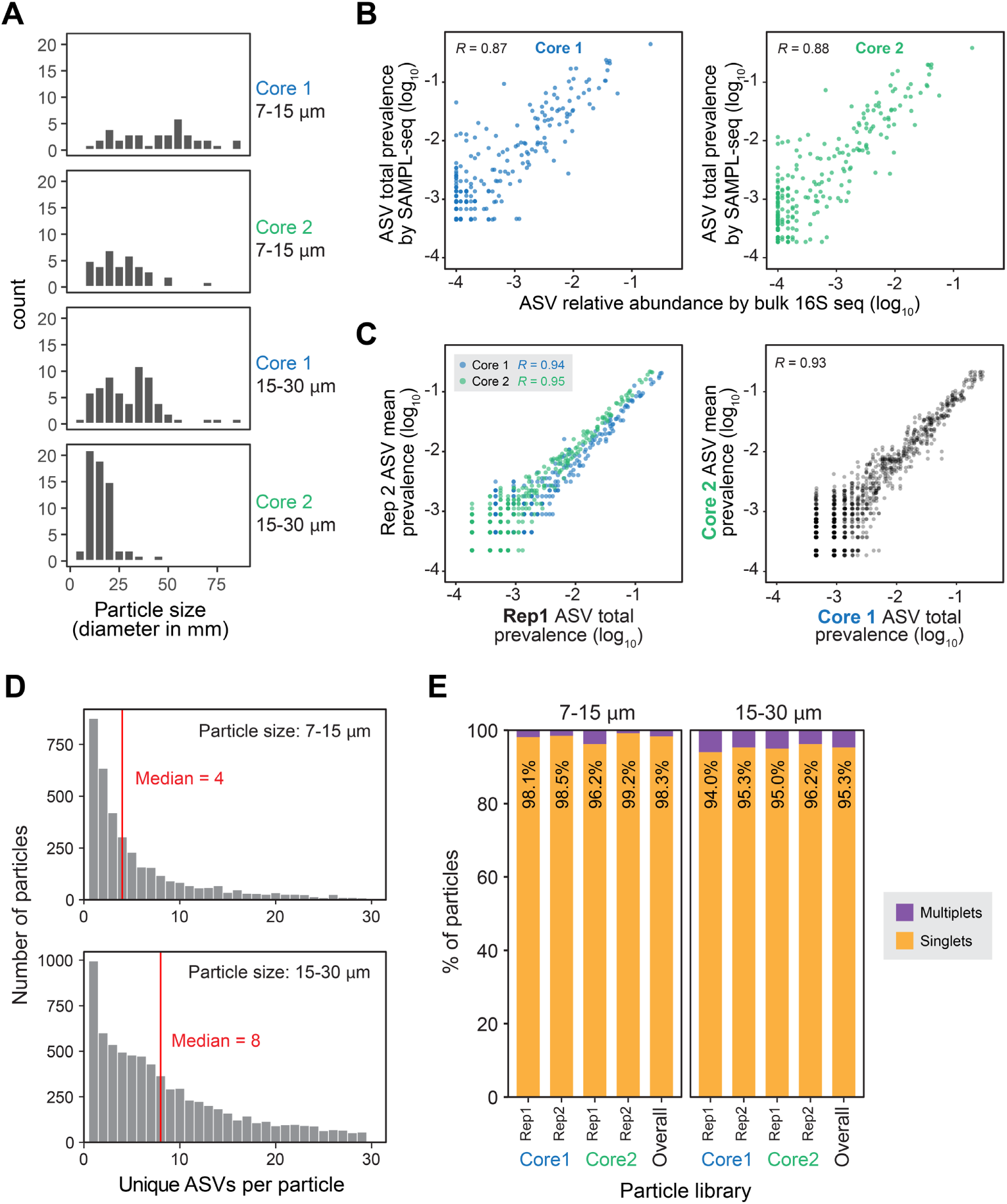
Homogenized fecal mixing experiment (M1). **(A)** Histograms of particle sizes for the replicates. **(B,C)** Correlation of ASV prevalence by SAMPLE-seq and ASV relative abundance by bulk 16S sequencing for Core 1 **(B)** and Core 2 **(C)**. SAMPL-seq abundances are averaged between replicates (excluding Spike-in)**. (D)** Histogram of the ASV per particle distribution by size (excluding Spike-in). **(E)** Barplot of the singlet rate of each replicate, grouped by particle size.

**Supplementary Figure 3.**
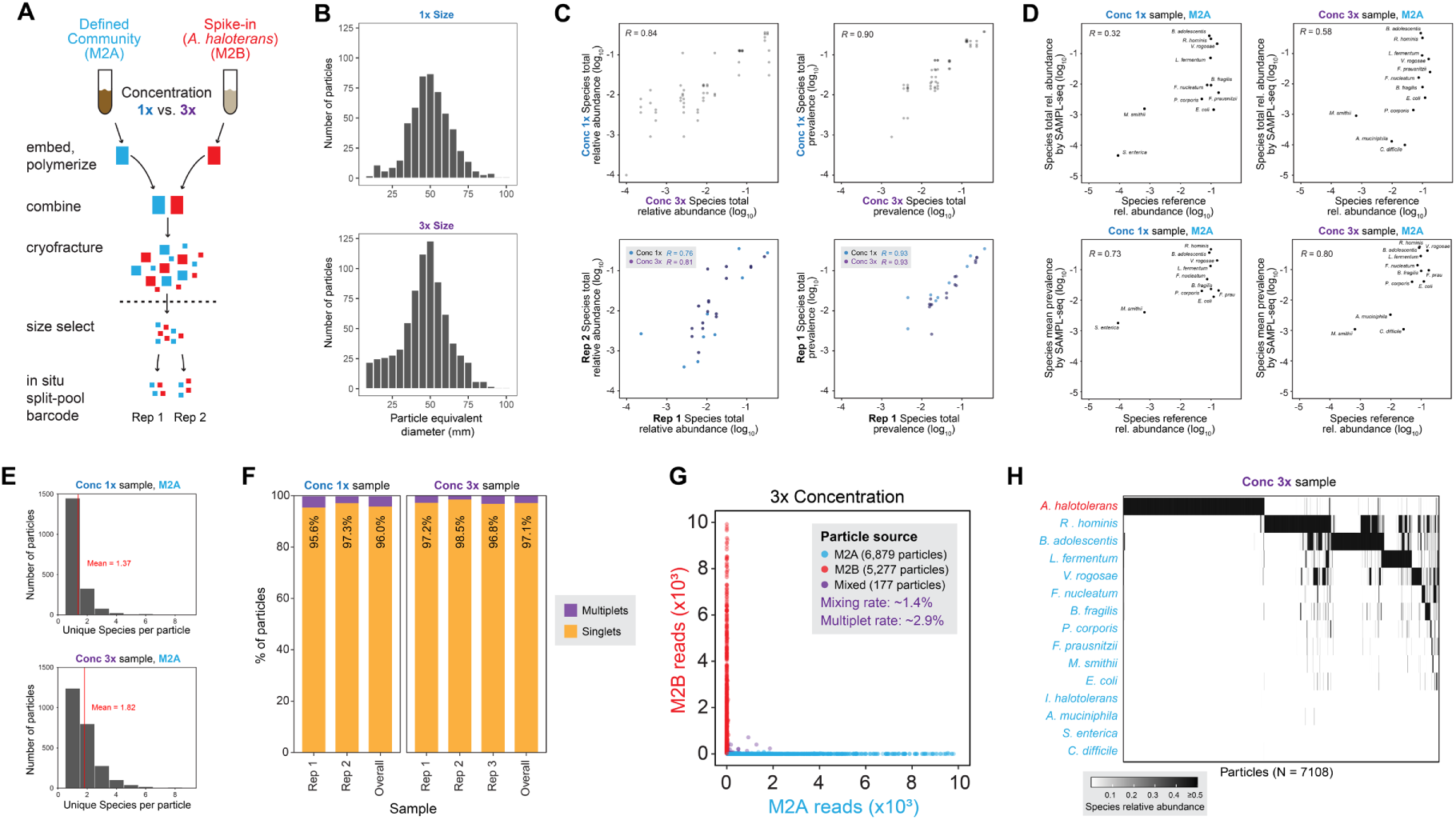
Defined community mixing experiment (M2). **(A)** An outline of the process for producing the fecal mixing library. Zymo Gut Microbial Standard and Zymo High Concentration Spike-in are separately embedded at equal cell ratios at 1x or 3x concentration replicates. They are then combined during the cryofracturing step, and are then size sorted, amplified and sequenced in aliquots of 10,000 particles. **(B)** Histograms of particle sizes for the replicates. **(C)** Scatterplots of technical (amplification) and biological (concentration) replicates, using both the relative abundance based on summed reads, and ASV prevalence among particles, which is the percentage of particles an ASV is found (excluding the Spike-in). **(D)** Scatterplot of ASV relative abundance and prevalence compared to absolute reference provided by the manufacturer. SAMPL-seq abundances are averaged between replicates (excluding Spike-in). **(E)** Histograms of the ASV per particle distribution by concentration (excluding Spike-in). **(F)** Barplot of the multiplet rate of each replicate, grouped by concentration**. (G)** Plot showing mixing rates of two defined communities (M1A and M1B), with each colored dot corresponding to a classified particle. **(H)** Heatmap of particles clustered by Bray-Curtis similarity and the Ward’s method.

**Supplementary Figure 4.**
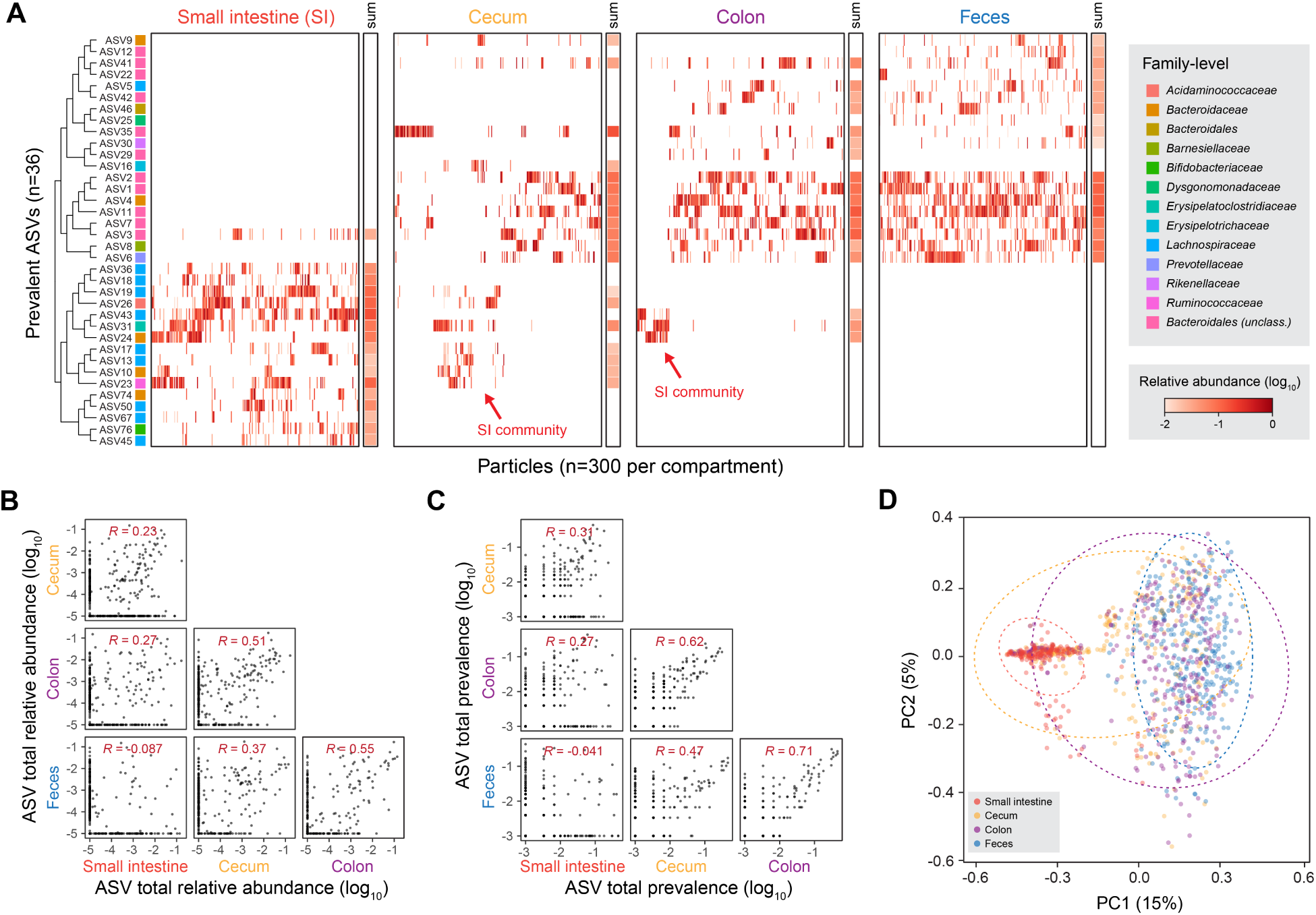
Correlation between ASVs from different murine gut compartments. **(A)** Clustered heatmap of prevalent mouse ASVs grouped by gut compartments, with summed abundances at the end of the row. ASVs are clustered by Jaccard overlap across the dataset. **(B,C)** Correlation between ASV relative abundance **(B)** and prevalence among particles **(C)** between mouse gut compartments. Colon and feces samples showed the highest correlation among samples. **(D)** Principal Coordinate Analysis (PCoA) plot of particles derived from different mouse gut compartments using Simpson distance, colored by gut compartments. Dashed circles correspond to the 95% confidence interval for each compartment using the multivariate t-distribution.

**Supplementary Figure 5.**
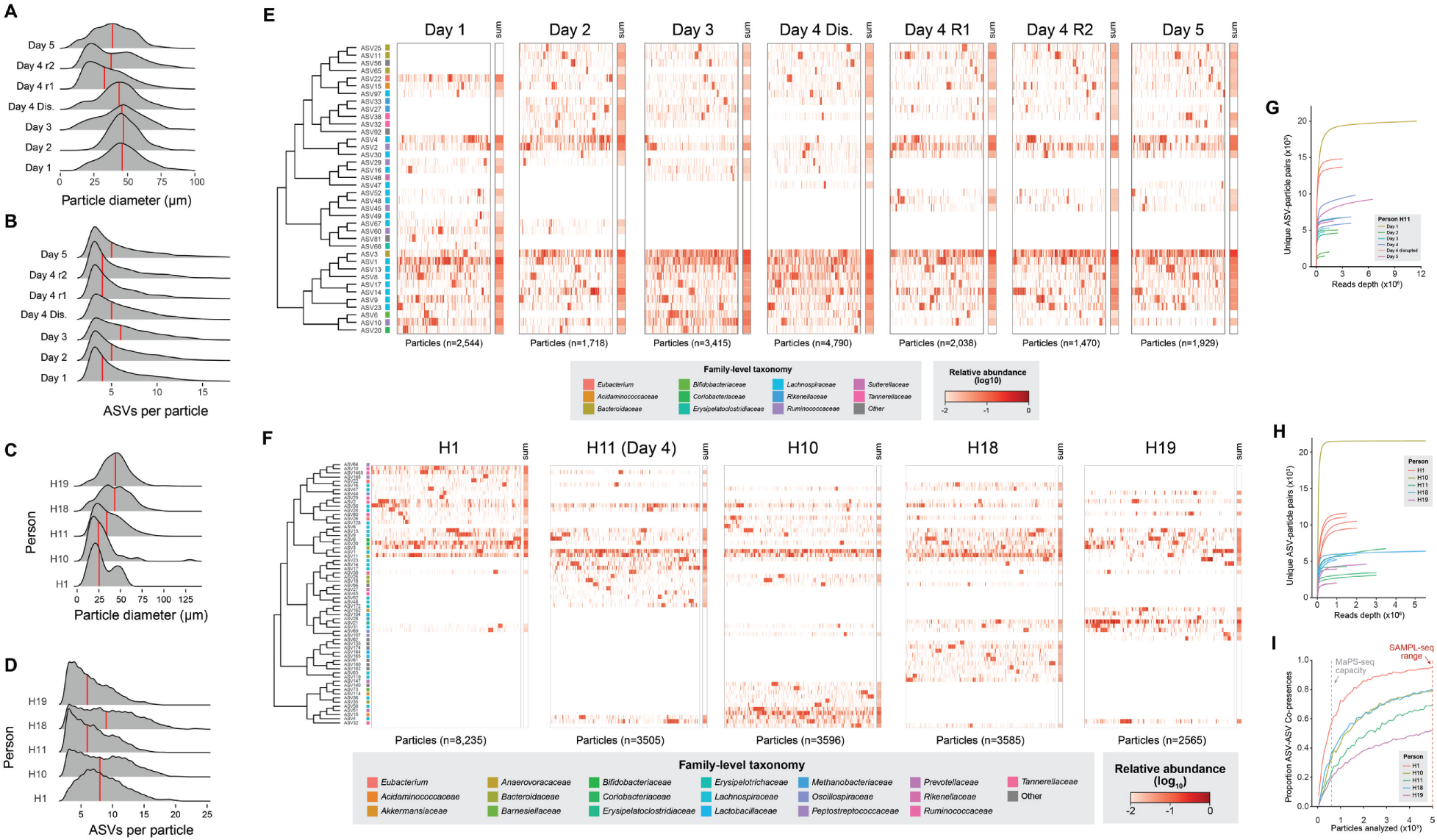
Particle-level data of the human gut microbiome from stool profiling. **(A,B)** Using filters for 20-40μm, distributions of particle sizes **(A)** and ASVs per particle **(B)** for longitudinal human stool samples from H11 are shown. **(C,D)** Using filters for 20-40μm, distributions of particle size **(C)** and ASVs per particle **(D)** for interpersonal samples are shown. **(E,F)** Particles from longitudinal **(E)** or interpersonal **(F)** are clustered within each day using the Simpson overlap, and ASVs are clustered using their Jaccard overlap across all days. **(G)** Rarefaction plot for longitudinal samples of unique ASV-particle pairs for prevalent ASVs (>1% prevalence in particles). **(H,I)** Rarefaction plots for interpersonal samples of unique ASV-particle pairs **(H)** for prevalent ASVs (>1% particle prevalence) or unique ASV-ASV co-presence **(I)** in a particle (observed >3 times).

**Supplementary Figure 6.**
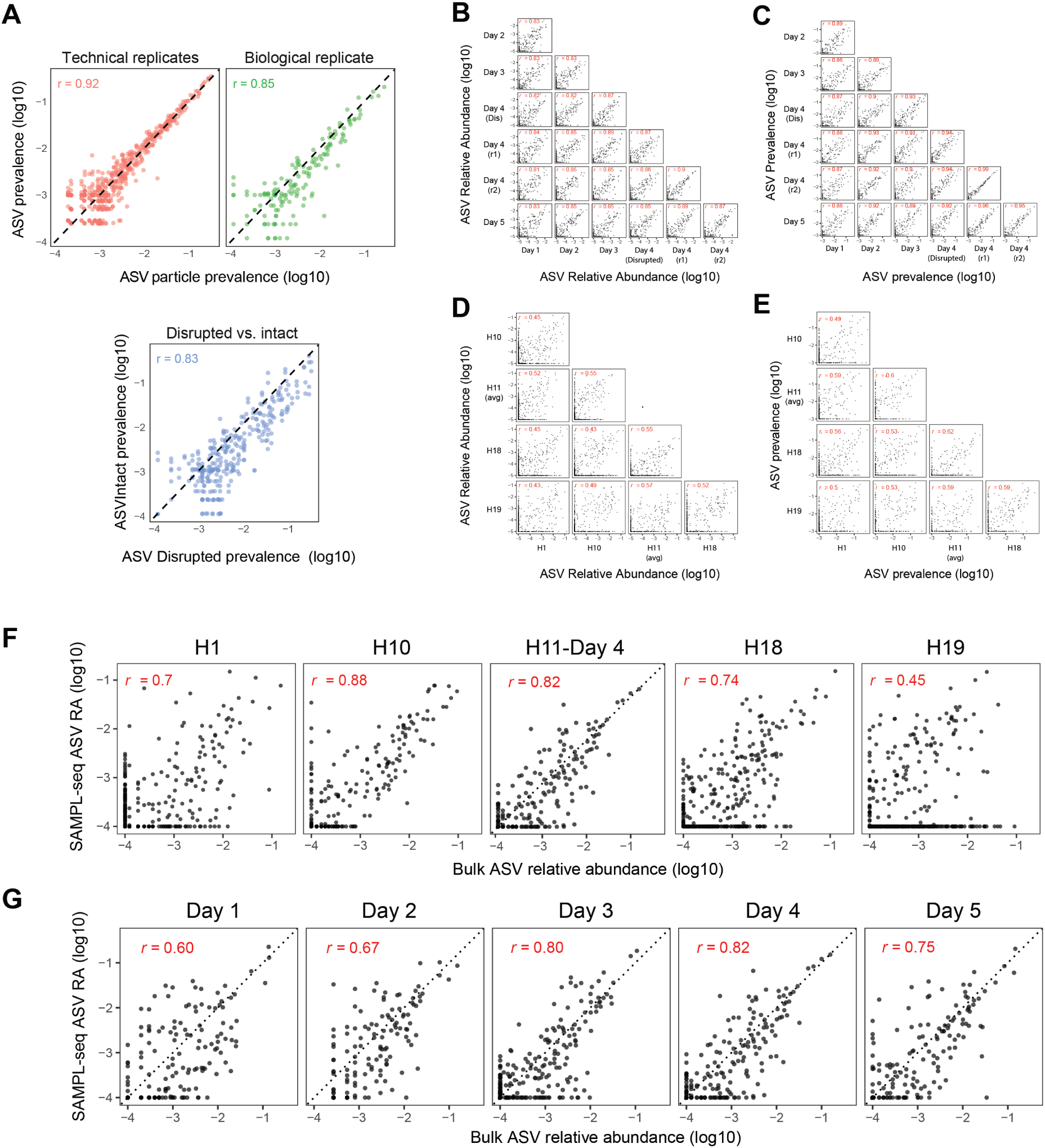
Technical validations of H11 Day 4 SAMPL-seq libraries. **(A)** Scatterplots of amplification (technical) replicates showed high correlation. Correlation between spatial (biological) replicates also showed high correlation. Homogenized sample showed high correlation, but increased particle prevalence relative to intact libraries. **(B,C)** Correlation of bulk ASV relative abundance **(B)** or ASV prevalence **(C)** between longitudinal SAMPL-seq libraries. **(D,E)** Correlation of bulk ASV relative abundance **(D)** or ASV prevalence **(E)** between interpersonal SAMPL-seq libraries**. (F,G)** Correlation of ASV relative abundance between interpersonal **(F)** or H11 longitudinal **(G)** samples with their corresponding bulk measurements.

**Supplementary Figure 7.**
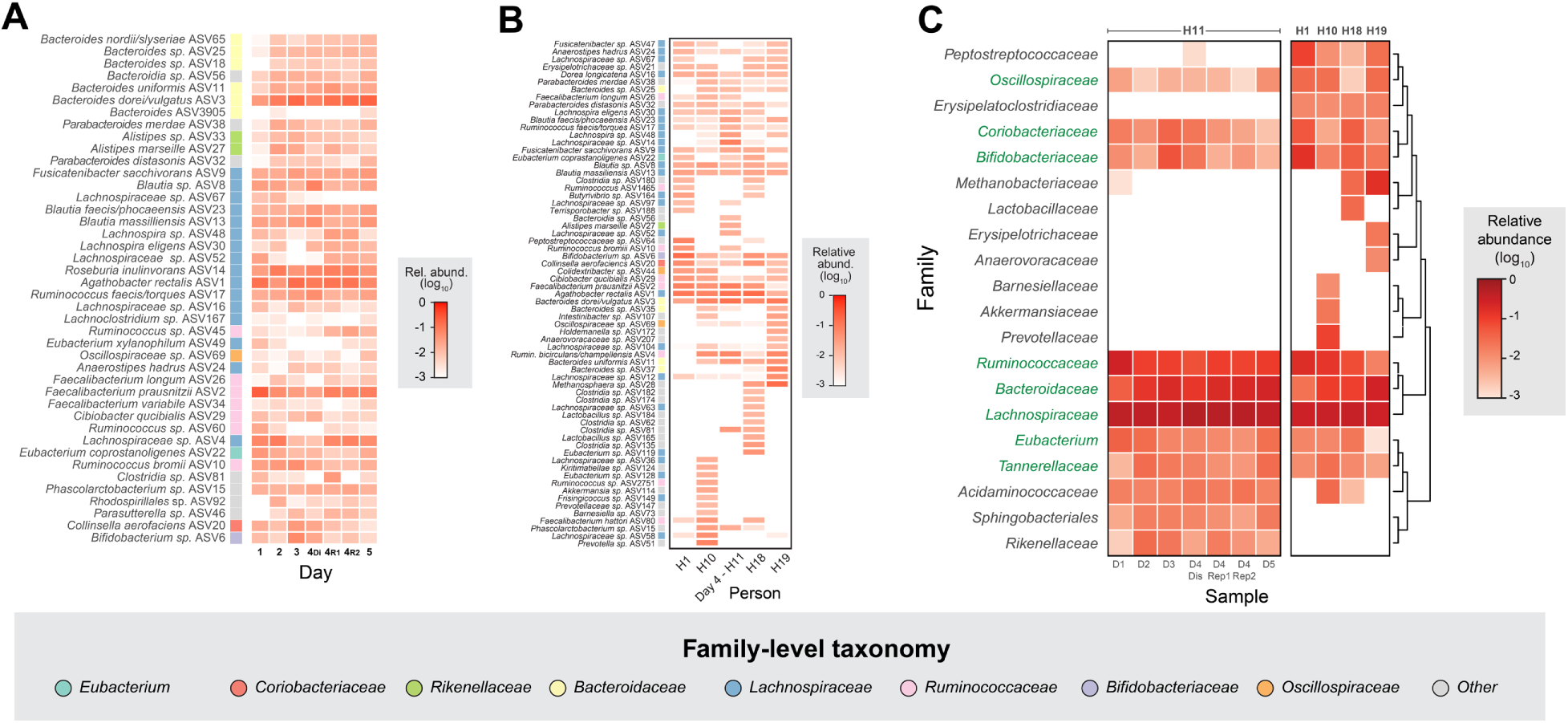
Large scale ASV compositional patterns. **(A)** Heatmap of overall ASV abundance in the dataset of prevalent ASVs (>1%), clustered by Jaccard overlap. **(B)** Heatmap of overall ASV abundance of prevalent ASVs (>1%) across 5 humans (H1, H10, H11, H18, H19), clustered by Bray-Curtis distance. **(C)** Heatmap of family-level relative abundance of human fecal samples. Families are clustered using the Jaccard overlap, and families conserved across all individuals are indicated in green.

**Supplementary Figure 8.**
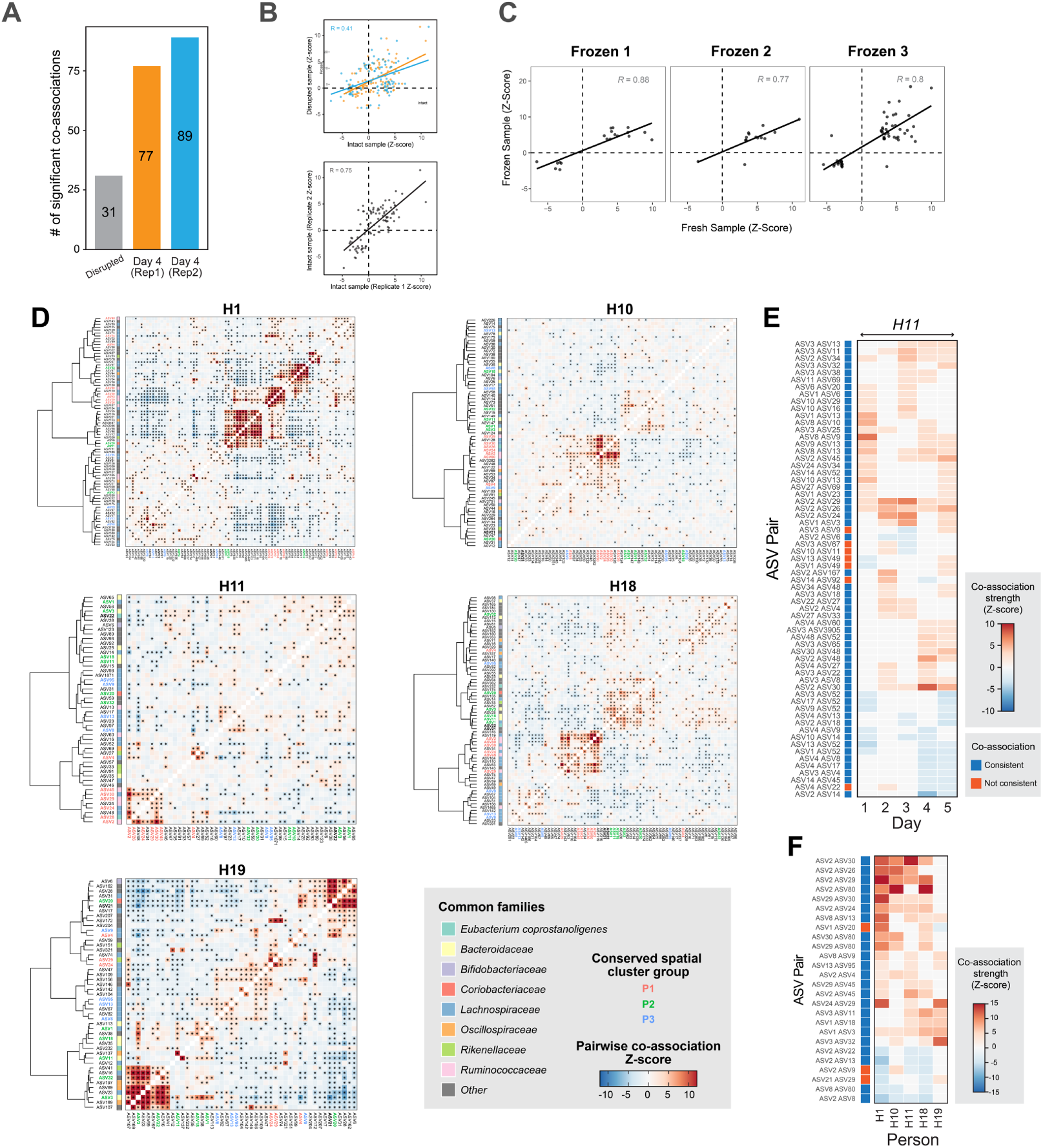
Pairwise ASV colocalization analysis. **(A)** Barplot of the number of significant ASV pairs found in each sample. **(B)** Scatterplot between ASV-pair Z-Scores between intact and disrupted samples. **(C)** Scatterplots of ASV-Pair Z-scores between fresh and frozen samples. **(D)** Pairwise ASV spatial associations in five people. Each heatmap shows all statistically significant spatial associations between pairs of ASVs for each individual (H1, H10, H11, H18, H19). Colors in the heatmap correspond to Z-scores and stars correspond to statistical significance (p<0.05 BH FDR Corrected). ASVs are labeled in 3 possible colors (red, green, blue) if they belong to a conserved spatial cluster group (P1, P2, P3) found across 3 or more individuals. Common taxonomic families are labeled next to each ASV label on the y-axis. **(E)** Heatmap of significant co-associations found on 2 or more days in H11. **(F)** Heatmap of significant co-associations found in 3 or more people.

**Supplementary Figure 9.**
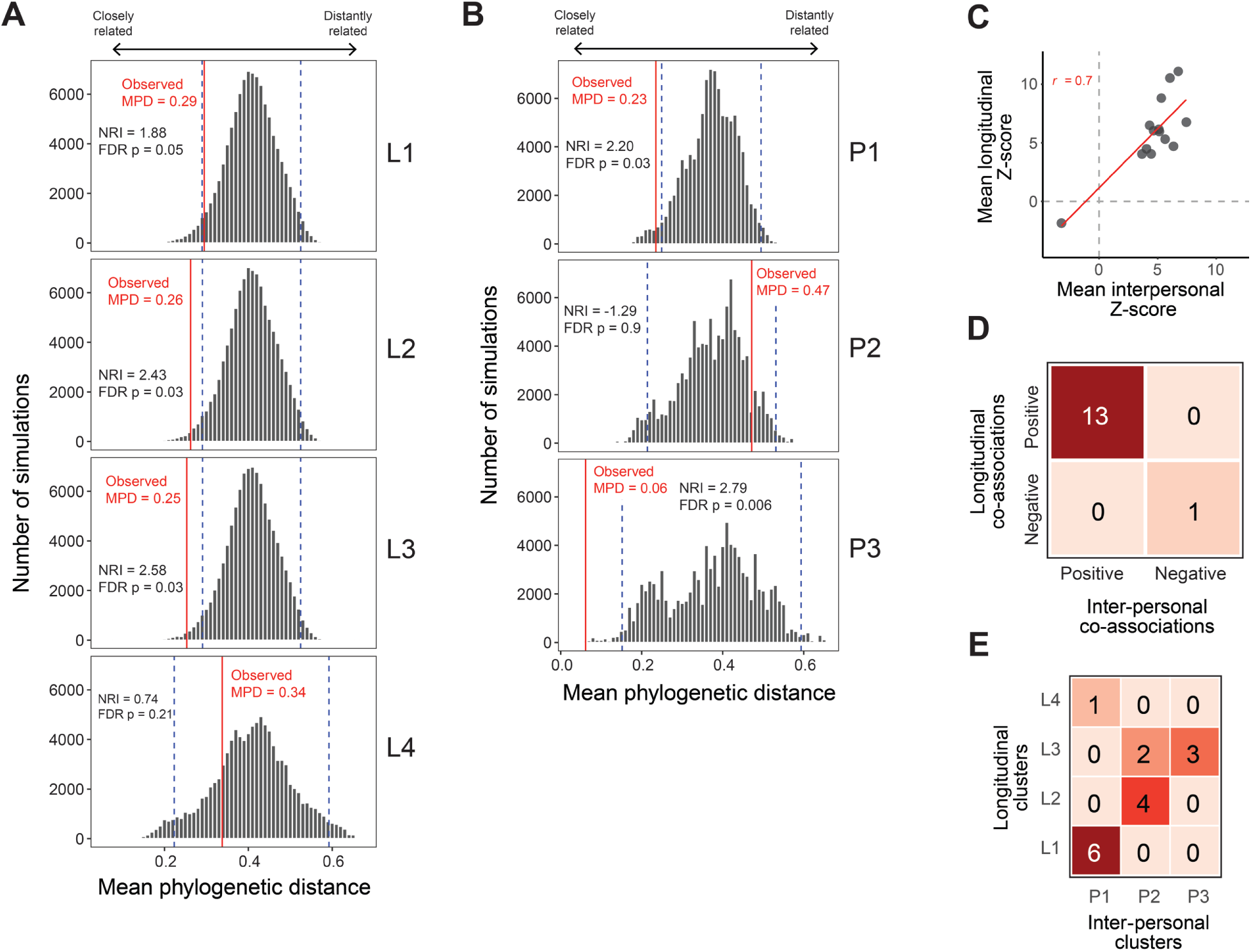
Phylogenetic Distance Distributions and Relationships Between Clusters. Histograms of simulated MPD distributions for L1-L4 **(A)** or P1-P3 **(B)** spatial hubs. The red line indicates the observed MPD in the cluster, while blue dashed lines indicate the 95% confidence interval around the mean of simulations. **(C)** Scatterplot of Z-score for associations found in both for longitudinal and interpersonal samples with the corresponding correlation**. (D)** Contingency table of the sign of longitudinal versus interpersonal associations. **(E)** Contingency table of ASV presence across the clusters. Chi-squared test of independence (p = 0.001).

**Supplementary Figure 10.**
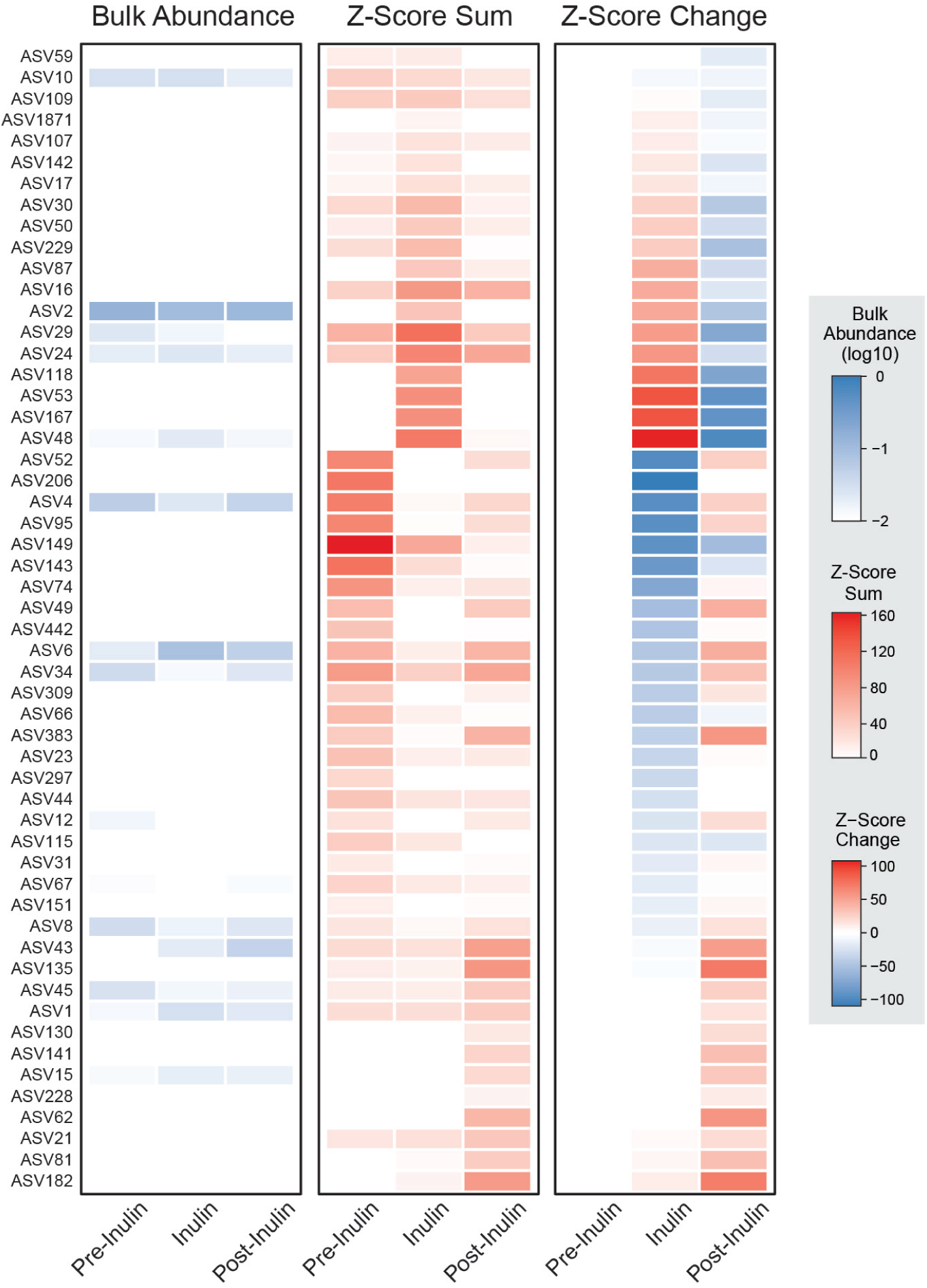
Inulin supplementation. Heatmaps of bulk relative abundance, Z-score sum, and change in total Z-score for ASVs that had a total Z-score change >10 over the course of inulin supplementation.

