## Supplementary Code for "SAMPL-seq reveals micron-scale spatial hubs in the human gut microbiome": Barcode Correction.R

```
# load libraries and set default plotting theme
library(data.table)
library(ggplot2)
library(stringr)
theme_set(theme_bw())

#################
melteddf2 <- fread("melted_otu_table.csv")

# take out the sample name
print("Extracting Sample Names")
melteddf2[,Sample := str_extract(Particle,"^.+(?=rbc)")]

# fix the samples (add bulk if present)
##### replace with your own filters
##### indicate whether a sample is a bulk sample or not
melteddf2[Bulk == "Particle"]

# remove bulk samples from BC correction
settocorrect <- melteddf2[Bulk == "Particle"]
unique(melteddf2$Bulk)

################
### Barcode extraction and analysis
# take out the individual barcodes from the particle name
print("Extracting Barcodes")
settocorrect[,Barcode1 := str_extract(Particle,"(?<=rbc).+")]
settocorrect[,ReadNum := str_extract(Barcode1,"_[0-9]+")]
settocorrect[,Barcode1 := str_remove(Barcode1,"_[0-9]+")]
settocorrect[,Barcode3 := str_extract(Barcode1,".{8}$")]
settocorrect[,Barcode1 := str_remove(Barcode1,".{8}$")]
settocorrect[,Barcode2 := str_extract(Barcode1,".{8}$")]
settocorrect[,Barcode1 := str_remove(Barcode1,".{8}$")]

# see how many unique barcodes we have
length(unique(settocorrect$Barcode1))
length(unique(settocorrect$Barcode2))
length(unique(settocorrect$Barcode3))

# read in our barcodes and assign them a barcode number
BC1 <- data.table(t(fread("../Barcodes/BC1.csv",header=F)))
BC1$BC1Number <- 1:96

BC1_7BP <- BC1[str_length(V1) == 7]
BC1_8BP <- BC1[str_length(V1) == 8]
BC1_9BP <- BC1[str_length(V1) == 9]

BC2 <- data.table(t(fread("../Barcodes/BC2.csv",header=F)))
BC2$BC2Number <- 1:96

BC3 <- data.table(t(fread("../Barcodes/BC3.csv",header=F)))
BC3$BC3Number <- 1:96

#############
### check what proportion of barcodes are error free
OverallCorrectionStatistics <- data.frame(rbind(table(settocorrect$Barcode1 %in% BC1$V1)/length(settocorrect$Barcode1),
      table(settocorrect$Barcode2 %in% BC2$V1)/length(settocorrect$Barcode2),
      table(settocorrect$Barcode3 %in% BC3$V1)/length(settocorrect$Barcode3)))

names(OverallCorrectionStatistics) <- c("BarcodesWithErrors","ErrorFreeBarcodes")
OverallCorrectionStatistics$BCRound <- c("BC1","BC2","BC3")
dir.create("Barcodes")
fwrite(OverallCorrectionStatistics,paste("Barcodes/",Sys.Date(),"PerfectvsErrorFreeBarcodes.csv"))

############
### do the actual correction
library(DNABarcodes)

# our barcodes are 3-8 hamming distance apart, so we can correct up to 1 error guaranteed
# run code below to see for yourself
#DNABarcodes::analyse.barcodes(BC2$V1)

# separate the BC1 barcodes because the software can't handle it...
bc1_7bp <- settocorrect$Barcode1[str_length(settocorrect$Barcode1) == 7]
bc1_8bp <- settocorrect$Barcode1[str_length(settocorrect$Barcode1) == 8]
bc1_9bp <- settocorrect$Barcode1[str_length(settocorrect$Barcode1) == 9]

# do the correction for each length set
print("Calculating Hamming Distances for BC 1")
bc1corrected7 <- demultiplex(unique(bc1_7bp), BC1_7BP$V1, metric="hamming")
bc1corrected8 <- demultiplex(unique(bc1_8bp), BC1_8BP$V1, metric="hamming")
bc1corrected9 <- demultiplex(unique(bc1_9bp), BC1_9BP$V1, metric="hamming")

# put the three lengths together,
# take the unique barcodes and see how they compare to reference
# get rid of barcode if it has hamming distance >1, as it cannot be safely corrected
bc1corrected <- data.table(rbind(bc1corrected7,bc1corrected8,bc1corrected9))
names(bc1corrected) <- c("OriginalBC1","CorrectedBC1","Distance")
bc1corrected[bc1corrected$Distance > 1,CorrectedBC1:=  NA]

print("Calculating Hamming Distances for BC 2")
bc2corrected <- data.table(demultiplex(unique(settocorrect$Barcode2), BC2$V1, metric="hamming"))
names(bc2corrected) <- c("OriginalBC2","CorrectedBC2","Distance")
bc2corrected[bc2corrected$Distance > 1,CorrectedBC2 :=  NA]

print("Calculating Hamming Distances for BC 3")
bc3corrected <- data.table(demultiplex(unique(settocorrect$Barcode3), BC3$V1, metric="hamming"))
names(bc3corrected) <- c("OriginalBC3","CorrectedBC3","Distance")
bc3corrected[bc3corrected$Distance > 1, CorrectedBC3 := NA]

################
### get number of corrections relative to barcode set

# get the particle read and unique particles counts for BC1, add them to the corrected bc set data.table
Barcode1Count <- settocorrect[,.(ParticleCount = .N,ReadCount = sum(Count)),by=Barcode1]
bc1corrected$ParticleBCCount <- Barcode1Count$ParticleCount[match(bc1corrected$OriginalBC1,Barcode1Count$Barcode1)]
bc1corrected$ReadBCCount <- Barcode1Count$ReadCount[match(bc1corrected$OriginalBC1,Barcode1Count$Barcode1)]

# calculate the number of particles/number of reads assigned to each hamming distance
BC1DistanceStat <- bc1corrected[,.(BarcodeReadCounts=sum(ReadBCCount),BarcodeParticleCounts=sum(ParticleBCCount)),by=Distance]
BC1DistanceStat$ProportionofBarcodesByParticles <- BC1DistanceStat$BarcodeParticleCounts/sum(BC1DistanceStat$BarcodeParticleCounts)
BC1DistanceStat$ProportionofBarcodesByReads <- BC1DistanceStat$BarcodeReadCounts/sum(BC1DistanceStat$BarcodeReadCounts)

# label them so it can be added with all the others
BC1DistanceStat$BarcodeSet <- "Barcode 1"

# get the particle read and unique particles counts for BC2, add them to the corrected bc set data.table
Barcode2Count <- settocorrect[,.(ParticleCount = .N,ReadCount = sum(Count)),by=Barcode2]
bc2corrected$ParticleBCCount <- Barcode2Count$ParticleCount[match(bc2corrected$OriginalBC2,Barcode2Count$Barcode2)]
bc2corrected$ReadBCCount <- Barcode2Count$ReadCount[match(bc2corrected$OriginalBC2,Barcode2Count$Barcode2)]

# calculate the number of particles/number of reads assigned to each hamming distance
BC2DistanceStat <- bc2corrected[,.(BarcodeReadCounts=sum(ReadBCCount),BarcodeParticleCounts=sum(ParticleBCCount)),by=Distance]
BC2DistanceStat$ProportionofBarcodesByParticles <- BC2DistanceStat$BarcodeParticleCounts/sum(BC2DistanceStat$BarcodeParticleCounts)
BC2DistanceStat$ProportionofBarcodesByReads <- BC2DistanceStat$BarcodeReadCounts/sum(BC2DistanceStat$BarcodeReadCounts)

# label them so it can be added with all the others
BC2DistanceStat$BarcodeSet <- "Barcode 2"

# get the particle read and unique particles counts for BC3, add them to the corrected bc set data.table
Barcode3Count <- settocorrect[,.(ParticleCount = .N,ReadCount = sum(Count)),by=Barcode3]
bc3corrected$ParticleBCCount <- Barcode3Count$ParticleCount[match(bc3corrected$OriginalBC3,Barcode3Count$Barcode3)]
bc3corrected$ReadBCCount <- Barcode3Count$ReadCount[match(bc3corrected$OriginalBC3,Barcode3Count$Barcode3)]

# calculate the number of particles/number of reads assigned to each hamming distance
BC3DistanceStat <- bc3corrected[,.(BarcodeReadCounts=sum(ReadBCCount),BarcodeParticleCounts=sum(ParticleBCCount)),by=Distance]
BC3DistanceStat$ProportionofBarcodesByParticles <- BC3DistanceStat$BarcodeParticleCounts/sum(BC3DistanceStat$BarcodeParticleCounts)
BC3DistanceStat$ProportionofBarcodesByReads <- BC3DistanceStat$BarcodeReadCounts/sum(BC3DistanceStat$BarcodeReadCounts)

# label them so it can be added with all the others
BC3DistanceStat$BarcodeSet <- "Barcode 3"

# combine them and export
AllStatistics <- data.table(rbind(BC1DistanceStat,BC2DistanceStat,BC3DistanceStat))
names(AllStatistics)[1] <- "HammingDistance"
fwrite(AllStatistics[order(BarcodeSet, HammingDistance)],"Barcodes/BarcodeCorrectionByDistance.csv")

# melt for plotting and fix up
HammingPlotFrame <- melt(AllStatistics[,.(HammingDistance,ProportionofBarcodesByParticles ,ProportionofBarcodesByReads,BarcodeSet)],id.vars = c("HammingDistance","BarcodeSet"))
HammingPlotFrame[HammingDistance < 2,BarcodeType := "Correctable"]
HammingPlotFrame[HammingDistance >= 2,BarcodeType := "Uncorrectable"]
HammingPlotFrame$HammingDistance <- factor(HammingPlotFrame$HammingDistance,levels=c("0","1","2","3","4","5"))
HammingPlotFrame$variable <- str_remove(as.character(HammingPlotFrame$variable),"ProportionofBarcodesBy") 

# plot faceted by barcode set
HammingPlot <- ggplot(HammingPlotFrame,aes(variable,value,fill=HammingDistance)) +
  geom_col(position = "stack") +
  xlab("Relative to the Number of") +
  ylab("Proportion of Total") +
  scale_fill_brewer(type="seq",direction=-1) +
  labs(fill="Hamming Distance") +
  facet_wrap(~BarcodeSet)
HammingPlot

ggsave("Barcodes/Hamming Distance Barplot By BC Round.pdf",HammingPlot,width=7,height=5)

# change to correctable vs. non correctable
correctableplotframe <- HammingPlotFrame[,.(PercentTotal = 100*sum(value)),by=.(BarcodeSet,variable,BarcodeType)]
correctableplotframe[,BarcodeType := factor(BarcodeType,levels=c("Uncorrectable","Correctable"))]

correctableplotframe[,PercentValue := round(PercentTotal,1)]
correctableplotframe[,PercentLabel := paste(PercentValue,"%",sep="")]
correctableplotframe[BarcodeType == "Uncorrectable",LabelPosition := 100-PercentValue/2]
correctableplotframe[BarcodeType == "Correctable",LabelPosition := 50]

# plot it 
CorrectableHammingPlot <- ggplot(correctableplotframe[variable=="Reads"],aes(BarcodeSet,PercentTotal,fill=BarcodeType)) +
  geom_col(position = "stack") +
  xlab("Barcode") +
  ylab("Percent of Barcodes") +
  theme_bw() +
  geom_text(aes(label=PercentLabel,y=LabelPosition ),size=2) +
  labs(fill="Barcode Type") +
  scale_y_continuous(breaks = seq(0,100,20))
CorrectableHammingPlot

ggsave("Barcodes/Correctable Hamming Distance Barplot By BC Round.pdf",CorrectableHammingPlot,width=5,height=5)

# plot it
HammingPlot <- ggplot(HammingPlotFrame[variable=="Reads"],aes(BarcodeSet,value,fill=HammingDistance)) +
  geom_col(position = "stack") +
  xlab("Relative to the Number of") +
  ylab("Proportion of Total") +
  scale_fill_brewer(type="seq",direction=-1) +
  labs(fill="Number Errors\n(Hamming Distance)")+
  scale_y_continuous(breaks = seq(0,100,20)) 
HammingPlot

ggsave("Barcodes/Hamming Distance Barplot By BC Round only reads.pdf",HammingPlot,width=7,height=5)


# plot faceted by particles/reads
HammingPlot2 <- ggplot(HammingPlotFrame,aes(BarcodeSet,value,fill=HammingDistance)) +
  geom_col(position = "stack") +
  xlab("Relative to the Number of") +
  ylab("Proportion of Total") +
  scale_fill_brewer(type="seq",direction=-1) +
  labs(fill="Hamming Distance") +
  facet_wrap(~variable)
HammingPlot2

ggsave("Barcodes/Hamming Distance Barplot By Relative Type.pdf",HammingPlot2,width=7,height=5)


############
### get the corresponding particle numbers
bc1corrected$Barcode1Number <- BC1$BC1Number[match(bc1corrected$CorrectedBC1,BC1$V1)]
bc2corrected$Barcode2Number <- BC2$BC2Number[match(bc2corrected$CorrectedBC2,BC2$V1)]
bc3corrected$Barcode3Number <- BC3$BC3Number[match(bc3corrected$CorrectedBC3,BC3$V1)]

# add them to the datatable
print("Correcting Particle Barcodes")
settocorrect$Barcode1Number <- bc1corrected$Barcode1Number[match(settocorrect$Barcode1,bc1corrected$OriginalBC1)]
settocorrect$Barcode2Number <- bc2corrected$Barcode2Number[match(settocorrect$Barcode2,bc2corrected$OriginalBC2)]
settocorrect$Barcode3Number <- bc3corrected$Barcode3Number[match(settocorrect$Barcode3,bc3corrected$OriginalBC3)]

##############
### get the number of unmatched barcodes per particle (this works because TRUE gets the value of 1 in R) ###
settocorrect[,UncorrectableBarcodes := (is.na(settocorrect$Barcode1Number) + is.na(settocorrect$Barcode2Number) + is.na(settocorrect$Barcode3Number))]
settocorrect[,CorrectlyAssignedBarcode := UncorrectableBarcodes <1]

bcprop <- settocorrect[,.(BarcodeReadCount = sum(Count)),by=.(CorrectlyAssignedBarcode)]
bcprop[,Dummy := "Yes"]
bcprop[,BarcodeAssigned := "No"]
bcprop[CorrectlyAssignedBarcode == TRUE,BarcodeAssigned := "Yes"]

bcprop[,BarcodePercent := round(BarcodeReadCount/sum(BarcodeReadCount)*100,2)]

totalpassingbarcode <- ggplot(bcprop,aes(x=Dummy ,y=BarcodePercent,fill=BarcodeAssigned)) +
  geom_col(position = "stack") +
  geom_text(aes(label=round(BarcodePercent,3)),color="black",size=5,position=position_stack(vjust=0.5)) +
  ylab("Percent of Reads") +
  theme(axis.text.x=element_blank(), 
        axis.ticks.x=element_blank()) +
  xlab("Overall")+
  scale_y_continuous(breaks = seq(0,100,20))

ggsave("Barcodes/Total Reads With Corrected and Assigned Barcodes.pdf",width=3,height=4)

NumberOfUncorrectedBCsPerParticle <- table(is.na(settocorrect$Barcode1Number) + is.na(settocorrect$Barcode2Number) + is.na(settocorrect$Barcode3Number))
fwrite(data.table(NumberOfUncorrectedBCsPerParticle),"Barcodes/NumberOfUncorrectedBCsPerParticle.csv")

##############
### see how many are ambiguous and export
barcodesummary <- data.table(rbind(table(is.na(settocorrect$Barcode1Number))/(length(settocorrect$Barcode1Number)),
      table(is.na(settocorrect$Barcode2Number))/(length(settocorrect$Barcode2Number)),
      table(is.na(settocorrect$Barcode3Number))/(length(settocorrect$Barcode3Number))))

# name them for output
names(barcodesummary) <- c("Corrected","Uncorrected")
barcodesummary$BC <- c("FirstBC","SecondBC","ThirdBC")
fwrite(barcodesummary,"Barcodes/BarcodeCorrectionFinalSummary.csv")

# making a plot
ambiguousplotframe <- melt(barcodesummary,id.vars = "BC")

AmbiguousPlot <- ggplot(ambiguousplotframe,aes(BC,value,fill=variable)) +
  geom_col(position = "stack") +
  xlab("Barcode Round") +
  ylab("Proportion of Particles")+
  labs(fill="Particle Barcode")

ggsave("Barcodes/Ambiguous Particle Barplot.pdf",AmbiguousPlot,width=5,height=5)

##############
### get rid of particles with ambiguous barcodes
finaldf <- settocorrect[complete.cases(settocorrect)]

##############
### get filtering statistics

# get pre/post data and merge
NumberParticlesBySample <- settocorrect[,.(ReadsPreCorrection=sum(Count),NumberParticlesPreBCCorrection = length(unique(Particle))),by=.(Sample)]
NumberParticlesAfterCorrection <- finaldf[,.(ReadsPostCorrection=sum(Count),NumberParticlesPostBCCorrection = length(unique(Particle))),by=.(Sample)]
FilteringStatistics <- merge(NumberParticlesBySample,NumberParticlesAfterCorrection,by="Sample")

# calculate percentages and write in nice order
FilteringStatistics$ProportionParticlesRemaining <-round(FilteringStatistics$NumberParticlesPostBCCorrection/FilteringStatistics$NumberParticlesPreBCCorrection,3)
FilteringStatistics$ProportionReadsRemaining <- round(FilteringStatistics$ReadsPostCorrection/FilteringStatistics$ReadsPreCorrection,3)
fwrite(FilteringStatistics[,c(1,2,4,7,3,5,6)],"Barcodes/Sample Filtering Statistics.csv")

# melt and plot
SampleStatisticsFrame <- melt(FilteringStatistics[,.(Sample ,ProportionParticlesRemaining,ProportionReadsRemaining)],id.vars = "Sample")
SampleStatisticsFrame$variable <- str_remove(SampleStatisticsFrame$variable,"Proportion")
SampleStatisticsFrame$variable <- str_replace(SampleStatisticsFrame$variable,"Remaining"," Remaining")

SampleStatistics <- ggplot(SampleStatisticsFrame,aes(Sample,value,fill=variable)) +
  geom_col(position = "dodge") +
  xlab("Sample") +
  ylab("Proportion")+
  labs(fill="Final Result") +
  theme(axis.text.x = element_text(angle = 270))

ggsave("Barcodes/Sample Filtering Statistics Boxplot.pdf",SampleStatistics,width=15,height=6)

#

################
### get number of corrections relative to barcode set

# get the particle read and unique particles counts for BC1, add them to the corrected bc set data.table
Barcode1SampleCount <- settocorrect[,.(ParticleCount = .N,ReadCount = sum(Count)),by=.(Barcode1,Sample)]
Barcode1SampleCount$Distance <- bc1corrected$Distance[match(Barcode1SampleCount$Barcode1,bc1corrected$OriginalBC1)]
Barcode1SampleCount$BarcodeRound <- "Barcode1"
Barcode1SampleCount$Barcode1 <- NULL

Barcode2SampleCount <- settocorrect[,.(ParticleCount = .N,ReadCount = sum(Count)),by=.(Barcode2,Sample)]
Barcode2SampleCount$Distance <- bc2corrected$Distance[match(Barcode2SampleCount$Barcode2,bc2corrected$OriginalBC2)]
Barcode2SampleCount$BarcodeRound <- "Barcode2"
Barcode2SampleCount$Barcode2 <- NULL

Barcode3SampleCount <- settocorrect[,.(ParticleCount = .N,ReadCount = sum(Count)),by=.(Barcode3,Sample)]
Barcode3SampleCount$Distance <- bc3corrected$Distance[match(Barcode3SampleCount$Barcode3,bc3corrected$OriginalBC3)]
Barcode3SampleCount$BarcodeRound <- "Barcode3"
Barcode3SampleCount$Barcode3 <- NULL

AllBCs <- rbind(Barcode1SampleCount,Barcode2SampleCount,Barcode3SampleCount)

SampleBCDistancePlot <- AllBCs[,.(NumberParticlesDistance = sum(ParticleCount),
                                  NumberReadsDistance = sum(ReadCount)),
                               by=.(Distance,BarcodeRound,Sample)]
is.data.table(SampleBCDistancePlot)
SampleBCDistancePlot <- SampleBCDistancePlot[ , SampleNumberParticles:= sum(NumberParticlesDistance),
                               by=.(BarcodeRound,Sample)]

SampleBCDistancePlot <- SampleBCDistancePlot[ , SampleNumberReads:= sum(NumberReadsDistance),
                                              by=.(BarcodeRound,Sample)]
SampleBCDistancePlot$ProportionParticle <- SampleBCDistancePlot$NumberParticlesDistance/SampleBCDistancePlot$SampleNumberParticles
SampleBCDistancePlot$ProportionReads <- SampleBCDistancePlot$NumberReadsDistance/SampleBCDistancePlot$SampleNumberReads

# combine them and export
fwrite(SampleBCDistancePlot[order(Sample,BarcodeRound, Distance)],"Barcodes/BarcodeCorrectionByDistanceSample.csv")

# melt for plotting and fix up
HammingPlotFrame <- melt(SampleBCDistancePlot[,.(Distance,ProportionParticle ,ProportionReads,BarcodeRound,Sample)],id.vars = c("Distance","BarcodeRound","Sample"))
HammingPlotFrame$Distance <- factor(HammingPlotFrame$Distance,levels=c("0","1","2","3","4","5"))
HammingPlotFrame$variable <- str_remove(as.character(HammingPlotFrame$variable),"Proportion") 

# plot faceted by barcode set
HammingPlot <- ggplot(HammingPlotFrame,aes(variable,value,fill=Distance)) +
  geom_col(position = "stack") +
  xlab("Relative to the Number of") +
  ylab("Proportion of Total") +
  scale_fill_brewer(type="seq",direction=-1) +
  labs(fill="Hamming Distance") +
  facet_wrap(Sample~BarcodeRound)

ggsave("Barcodes/Hamming plot by Sample.pdf",HammingPlot,width=40,height=40)


##########
### create corrected particle IDs
print("Creating Final data.table")

finaldf$CorrectedParticleID <- paste(finaldf$Sample,"bcnum",finaldf$Barcode1Number,"_",finaldf$Barcode2Number,"_",finaldf$Barcode3Number,sep="")

# combine with bulk samples and correct names
finaldfbulk <- rbind(finaldf,melteddf2[Bulk == "Bulk"],fill=T)
finaldfbulk$CorrectedParticleID[finaldfbulk$Bulk == "Bulk"] <- finaldfbulk$Particle[finaldfbulk$Bulk == "Bulk"]

# resum by corrected particle IDs
CorrectedDataframe <- finaldfbulk[, sum(Count), by = .(CorrectedParticleID,OTU,Sample)]
print("Unique Sample Names:")
print(unique(CorrectedDataframe$Sample))

# rename
names(CorrectedDataframe) <- c("Particle","OTU","Sample","Count")
print("Writing Final data.table")
fwrite(CorrectedDataframe,"CorrectedDataFrame.csv")
```
