## Supplementary Code for "SAMPL-seq reveals micron-scale spatial hubs in the human gut microbiome": Barcoding Processing.bash

```
# housekeeping

# this script requires a bin of software/resources, including usearch/vsearch and the primer sequences in the local directory
# the primer sequences are included on github, but I cannot distribute usearch without a license

# we put all fastqs to process in the fastq folder, everything else should work from there

#### setup your new machine with all the requisite software ####
# I use an ubuntu 18.04 machine

# make sure repositories are up to date
sudo apt-get update
sudo apt-get upgrade

# install parallel
sudo apt-get install parallel 

# get necessary software from conda
conda install -c conda-forge -c bioconda ultraplex
conda install -c bioconda seqkit
conda install -c bioconda vsearch
conda install -c conda-forge r-base 

# USEARCH is not included here but can be found at https://www.drive5.com/usearch/

#### Processing Samples ####

#unzip fastqs for usearch
unpigz ./fastq/*

# filter reads from individual samples
foo () {
    local FILENAME=$1
    OUTFILE="$(echo ${FILENAME} | sed 's/_.*$//')_filtered.fastq";
    ./bin/usearch10 -fastq_filter "${FILENAME}" -fastq_maxee 1.0 -fastq_minlen 150 -relabel @ -fastqout "${OUTFILE}" -threads 2
}

for run in find ./fastq/*R1* -type f; do foo "$run" & done


#remove the original fastqs
rm ./fastq/*_R1_001.fastq.gz

# zip filtered fastqs for ultraplex
pigz ./fastq/*_filtered.fastq

#### extracting unique particles from barcodes MASSIVELY FASTER NOWWWWW ####
#unpacking sequencing files
find ./fastq/*_filtered.fastq.gz -type f | while read FILENAME ; do
    OUTFILE="$(echo ${FILENAME} | sed 's/_.*$//')_filtered_particleid.fastq.gz";
	
	#ultraplex for 7,8,9 bp leading barcodes (it can't do all at once...)
	# this could be solved by padding those with 7 and 8 bp with pads, but that is probably slower
	# minimum length 86bp, 32 threads, and minimum qscore 20 
	ultraplex -i "${FILENAME}" -b ./bin/BarcodeCSV7leadingfull.csv -l 86 -inm -t 32 -dbr -d fastq/first7bp -q 20
	ultraplex -i "${FILENAME}" -b ./bin/BarcodeCSV8leadingfull.csv -l 86 -inm -t 32 -dbr -d fastq/first8bp -q 20
	ultraplex -i "${FILENAME}" -b ./bin/BarcodeCSV9leadingfull.csv -l 86 -inm -t 32 -dbr -d fastq/first9bp -q 20

	#combine them
	echo "Combining Files"
	cat ./fastq/*first*bp/ultraplex*.fastq.gz > ./fastq/AllLengths.fastq.gz
	
	# strip 16S primer (the degenerate bases don't play well with seqkit)
	echo "Removing Primers"
	seqkit amplicon  ./fastq/AllLengths.fastq.gz -F YCAGCMGCCGCGGTAA -r 1:240 -f  -j 32 -o "${OUTFILE}";
	
	echo "Cleaning Up"
	# remove intermediates file (could be removed by piping this into the earlier command pipe?)
	rm ./fastq/AllLengths.fastq.gz
	
	# get rid of directory with the intermediate files
	rm ./fastq/first* -rf
	
	echo "Done"
done


#combine fastqs into one sample in new directory (fast)
mkdir otu

cat fastq/*_filtered_particleid.fastq.gz > otu/filt_all_out.fastq.gz

# fix the read names so usearch/vsearch will understand the samples (slow)
seqkit replace otu/filt_all_out.fastq.gz -p "\.[0-9]+rbc:" -r 'rbc' -o otu/filt_all_particleid.fastq.gz

# check the length
seqkit stats otu/filt_all_particleid.fastq.gz

# remove short reads (slow) mostly fixed by increasing ultraplex length (still got some weird short reads)
seqkit seq otu/filt_all_particleid.fastq.gz -m 69 -o otu/69bpfilt_all_particleid.fastq.gz -j 24 -g

# trim to 69bp (slow but not too bad)
seqkit subseq  otu/69bpfilt_all_particleid.fastq.gz  -r 1:69 -j 24 -o otu/final_filt_all_particleid.fastq.gz

#convert to fasta for VSEARCH clustering (fast)
seqkit fq2fa otu/final_filt_all_particleid.fastq.gz -o otu/filt_all.fa -j 24

# unzip your fastqs to make them useable with usearch
unpigz otu/final_filt_all_particleid.fastq.gz

#### OTU Clustering ####
# remove doubles (old way reduces otus by a lot ~70%)
./bin/usearch10 -fastq_filter otu/final_filt_all_particleid.fastq -fastq_maxee 0.1 -relabel @ -fastqout otu/filt_all_double.fq -threads 36

# get unique sequences
./bin/usearch10 -fastx_uniques otu/filt_all_double.fq -sizeout -relabel Uniq -fastaout otu/uniques.fa -threads 36

# cluster otus
./bin/usearch10 -unoise3 otu/uniques.fa -zotus otu/zotus.fa

# change Zotu to Otu
sed -i 's/Zotu/Otu/g' otu/zotus.fa

# make final files
mkdir ProcessedResults

# make udb (supposed to speed up search)
vsearch --makeudb_usearch ./otu/zotus.fa --output ./otu/zotus.udb

#do assignment with vsearch (much faster than usearch)
vsearch --usearch_global ./otu/filt_all.fa --db ./otu/zotus.udb --id 0.9 --otutabout ./ProcessedResults/otu_frequency_table.tsv --biomout ./ProcessedResults/otu_frequency_table.biom --threads 72

# add otus to ProcessedResults
cp otu/zotus.fa ProcessedResults/

#compress the frequency table, as it is often large
pigz ./ProcessedResults/otu_frequency_table.tsv
```
