## Supplementary Code for "SAMPL-seq reveals micron-scale spatial hubs in the human gut microbiome": Biom File Melting.R

```
library(data.table)

# read in the resulting otu table (biom is faster and has defined structure)
# http://biom-format.org/documentation/format_versions/biom-1.0.html
df <- jsonlite::fromJSON("otu_frequency_table.biom", flatten=TRUE)

# if matrix_type is "sparse", data = [[row, column, value]
melteddata <- data.table(df$data)

# An ORDERED list of obj describing the rows
rowmapping <- data.table(df$row)[,.(OTU =id)]
rowmapping$RowNum <- 0:(nrow(rowmapping)-1)

#An ORDERED list of obj  describing the columns
columnmapping <- data.table(df$columns)[,.(Particle = id)]
columnmapping$ColNum <- 0:(nrow(columnmapping)-1)

# match the row/column names and add to the DF
melteddata$OTU <- rowmapping$OTU[match(melteddata$V1,rowmapping$RowNum)]
melteddata$Particle <- columnmapping$Particle[match(melteddata$V2,columnmapping$ColNum)]

# subset for future things
formattedmelted <- melteddata[,.(Particle,OTU,Count=V3)]

# write it
fwrite(formattedmelted,"melted_otu_table.csv")
```
